# Intracellular pH dynamics regulates intestinal stem cell lineage specification

**DOI:** 10.1101/2021.10.28.466337

**Authors:** Yi Liu, Efren Reyes, David Castillo-Azofeifa, Ophir D. Klein, Diane L. Barber, Todd Nystul

**Author notes:** These authors contributed equally to this work (Efren Reyes, David Castillo-Azofeifa).

## Abstract

Intracellular pH (pHi) dynamics is increasingly recognized to regulate myriad cell behaviors, including proliferation, migration, differentiation, and transformation. Here we report a new finding that pHi dynamics also regulates adult stem cell lineage specification. In mouse small intestinal organoids, we identify a pHi gradient along the crypt axis, lower at the crypt base and higher toward the villus, and find that dissipating this gradient by inhibiting Na^+^-H^+^ exchanger 1 (NHE1) activity genetically or pharmacologically abolishes crypt budding. Using single-cell RNA sequencing and lineage tracing we demonstrate that pHi dynamics acts downstream of ATOH1, with increased pHi promoting differentiation toward the secretory lineage, while reduced pHi biases differentiation into the absorptive lineage. Consistent with these results, disrupting the pHi gradient blocks new Paneth cell differentiation. Paneth cells provide an essential WNT signal to ISCs in organoids, and we find that the loss of crypt budding with inhibiting NHE1 activity is rescued with exogenous WNTs. Our findings indicate that pHi dynamics is tightly regulated in the ISC lineage and that an increase in pHi is required for the specification of secretory lineage, including Paneth cell differentiation that contributes to crypt maintenance. These observations reveal a previously unreported role for pHi dynamics in cell fate decisions within an adult stem cell lineage.

## Introduction

Intracellular pH (pHi) dynamics is increasingly recognized as a key regulator of diverse cell behaviors, including epithelial to mesenchymal transition (Amith et al., 2016; Raja et al., 2020), transformation and dysplasia (Amith et al., 2016; Grillo-Hill et al., 2015; Liu et al., 2020; White et al., 2017), and differentiation (Benitez et al., 2019; Liu et al., 2020; Oginuma et al., 2020; Raja et al., 2020; Ulmschneider et al., 2016). We previously reported that increased pHi is required for differentiation of clonal mouse embryonic stem cells and *Drosophila* adult ovary follicle stem cells (Ulmschneider et al. 2016; Benitez et al. 2019). However, it remains unknown whether pHi dynamics regulates lineage specification of mammalian adult stem cells. To investigate this possibility, we used mouse small intestinal organoids as a model. Like the *Drosophila* follicle epithelium, the epithelium in these organoids is maintained by a pool of adult stem cells (intestinal stem cells, ISCs) that divide regularly during adult homeostasis to self-renew and produce daughter cells that first increase in number through a transit amplification stage and have the potential to differentiate along multiple different lineage trajectories (Beumer and Clevers, 2021; Castillo-Azofeifa et al., 2019; van Es et al., 2012a; Gehart and Clevers, 2019; Kurokawa et al., 2020; Sancho et al., 2015; Sato et al., 2009; Sprangers et al., 2021; Stanger et al., 2005; VanDussen and Samuelson, 2010; VanDussen et al., 2012).

To determine pHi dynamics in ISC behaviors we used the 3D organoids derived from the mouse duodenum. Intestinal organoids are a self-sustaining *in vitro* epithelial 3D model that recapitulates many features of *in vivo* mammalian intestinal epithelium, including self-renewing stem cells at the base of budding crypts, a distinct crypt-villus architecture, and differentiation and lineage specification of stem cell progeny (Beumer and Clevers, 2021; Boonekamp et al., 2020; Gehart and Clevers, 2019; Lukonin et al., 2020; Sachs et al., 2017; Serra et al., 2019; Sprangers et al., 2021). We identified a pHi gradient along the crypt axis in intestinal organoids that is lowest in the crypt base and increases up the crypt column, and found that the gradient is maintained by the activity of the plasma membrane H^+^ extruder NHE1. Single-cell RNA sequencing (scRNA-seq) revealed that disrupting the pHi gradient impairs specification of the secretory lineage downstream of the master regulator, ATOH1, which we confirmed with lineage tracing. Paneth cells are a major cell type in the secretory lineage, and we found that disrupting the pHi gradient resulted in loss of Paneth cell differentiation, which impairs the WNT circuit and crypt budding. In addition, the scRNA-seq revealed a biased stem cell fate decision toward the absorptive lineage and an increased *Clusterin* (*Clu*)^+^ revival subpopulation. Together, our results reveal a previously unreported role for pHi dynamics as a regulator of mammalian adult stem cell lineage specification.

## Results

### A pHi gradient via NHE1 activity in small intestinal organoids

To quantify the pHi of cells in the ISC lineage, we generated organoids stably expressing a genetically encoded pHi biosensor, mCherry-SEpHluorin (Benitez et al., 2019; Grillo-Hill et al., 2014, 2015; Ulmschneider et al., 2016). We acquired ratiometric (SEpHluorin/mCherry) images at days 1 and 3 of organoid growth (Figures 1A and 1B; Figure S1A) and calibrated fluorescent ratios to pHi by perfusing with nigericin-containing buffers of known pH values at the end of each imaging set (Grillo-Hill et al., 2014). We found that all cells have a similar pHi at day 1 of organoid growth but by day 3 a pHi gradient develops along the crypt axis (Figures 1B and 1C). We calibrated pHi values of ∼7.2 in the ISCs and Paneth cells, ∼7.4 in the progenitor cells on the crypt column and ∼7.5 in the crypt neck region (Figure 1C). Ratiometric time-lapse imaging also revealed a similar pHi gradient during crypt formation, with higher pHi in cells within the column and neck than in the crypt base (Figure 1D; Video S1). These differences in pHi raised the question of whether increased pHi regulates ISC differentiation.

**Figure 1.**
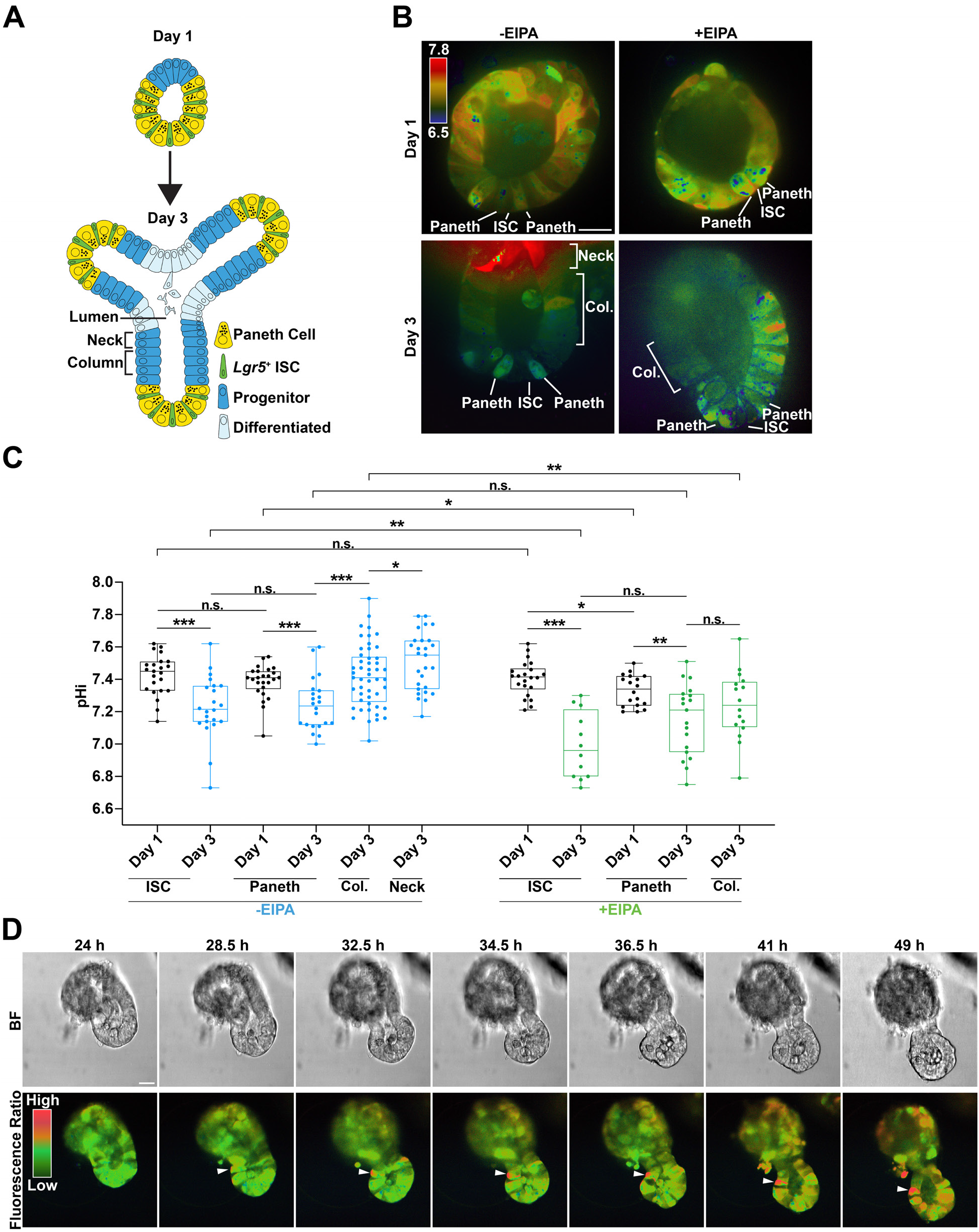
A pHi gradient develops in small intestinal organoid crypts. (**A**) Schematic representation of small intestinal organoids growth. (**B** to **D**) pHi in the intestinal crypt. (B) Representative ratiometric images of mCherry-SEpHluorin ratio (SEpHluorin/mCherry) in a WT organoid (control) and an WT organoid treated with 5 μM EIPA from day 1 to day 3. Images show organoids from a representative 3 preparations. Confocal images of individual fluorescent channels with single-cell resolution (see figure S1A). (C) Values of pHi of different crypt cell types determined by calibrating fluorescent ratios (n=3; Mann-Whitney test). A pHi gradient in the crypts of day 3 control (blue) is attenuated with EIPA (green). Box plots are minimum to maximum, the box shows 25th-75th percentiles, and the central line is the median. (D) Representative frames (7 of 51) from 15-hr time-lapse videos of crypt budding started 24 hr after seeding. A cell with a high pHi appears in the crypt neck and retained in the crypt column region during crypt growth (arrowhead). BF, bright-field view. Fluorescence Ratio, ratiometric view of mCherry-SEpHluorin. Time-lapse videos of pHi dynamics in crypt budding in movie S1.\**P* < 0.05, \*\**P* < 0.01, \*\*\**P* < 0.001. N.s., not statistically significant. All scale bars represent 20 μm.

To determine what regulates this pHi gradient, we tested the Na^+^-H^+^ exchanger 1 (NHE1), a nearly ubiquitously expressed resident plasma membrane protein that exchanges an influx of extracellular Na^+^ for an efflux of intracellular H^+^ that is a key regulator of pHi dynamics in mammalian cells (Cao et al., 2019; Putney et al., 2002; Zachos et al., 2005). Likewise, the ortholog of NHE1 in *Drosophila*, *dNhe2*, is necessary for a pHi gradient within the ovarian follicle stem cell lineage, with a lower pHi in follicle stem cells that progressively increases in pre-follicle and mature follicle cells (Ulmschneider et al., 2016). We found that intestinal organoids treated with 5 μM 5-(N-Ethyl-N-isopropyl)-amiloride (EIPA), a selective pharmacological inhibitor of NHE1 (Pedersen et al., 2007), had a significantly lower pHi in ISCs and adjacent non-Paneth cells compared with controls at day 3 (Figures 1B and 1C). In contrast, pHi remained unchanged in ISCs between controls and EIPA-treated organoids at day 1. Hence, inhibiting NHE1 activity reduced pHi in different crypt cells and disrupted the pHi gradient that was generated by day 3.

### NHE1 activity is necessary for crypt growth

To determine whether the pHi gradient along the crypt axis is functionally significant for organoid development, we disrupted the gradient by inhibiting NHE1 activity pharmacologically with EIPA and genetically with a doxycycline (Dox)-inducible CRISPR-Cas9 system (Methods). We used inducible gene silencing in organoids to limit compensatory changes reported to occur in other ion transport proteins in animals engineered to be null for pHi regulators (Christensen et al. 2013; Bachmann et al. 2004; Guan et al. 2006; Bailey et al. 2004; Nikolovska et al. 2022). Additionally, although NHE1 has dual functions as a plasma membrane ion transport protein and an actin filament anchor through binding the ERM proteins ezrin. radixin and moesin (Denker and Barber, 2002; Denker et al., 2000), current evidence indicates EIPA targets only ion transport activity by NHE1 with no reports of it affecting actin anchoring.

In control organoids, crypts began budding by day 1, then lengthened and enlarged by day 3 as mature buds. In contrast, inhibiting NHE1 activity with EIPA or by Dox-inducible CRISPR-Cas9 gene silencing (Figures 2A and 2C) significantly reduced the number of budded crypts (Figures 2B and 2D). When crypt buds did form in organoids treated with EIPA, they were typically not retained (Figure 2A; Video S2B). Additionally, budding resumed after removing of EIPA at day 3 (Figures 2E and 2F), indicating that organoids remain viable with EIPA treatment (Figure S1B) and that the inhibition of budding is reversible. To test whether loss of budding is due to the absence of crypt cells, we immunolabeled for the hyaluronic acid receptor CD44, a crypt marker expressed throughout the crypt cell population, including *Lgr5^+^* ISCs, Paneth cells, and crypt progenitors (Sumigray et al., 2018; Zeilstra et al., 2008, 2014). We found that CD44^+^ cell clusters were retained in organoids that lack NHE1 activity (Figure 2G). We also identified Paneth cells, as indicated by lysozyme (LYZ) immunolabeling, within the CD44^+^ population of cells (Figure 2G), suggesting that these CD44^+^ clusters are unbudded crypts. Taken together, these data indicate that loss of NHE1 activity disrupts crypt budding but does not eliminate crypt cell populations.

**Figure 2.**
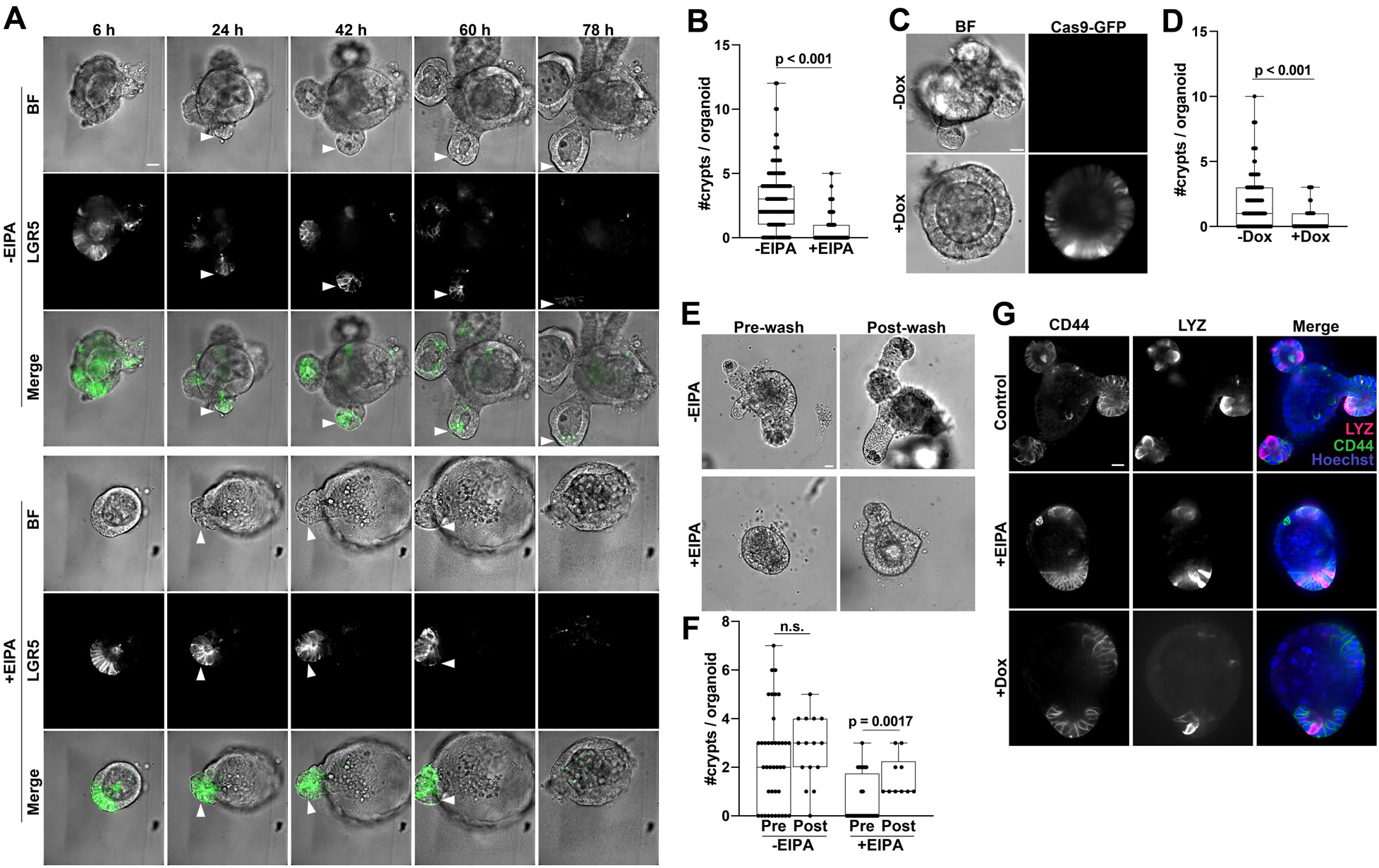
Loss of NHE1 activity attenuates intestinal crypt budding in organoids. (**A**) Representative frames (5 of 25) from 75-hr time-lapse microscopy (see videos S2A and S2B) showing crypt budding (arrowhead) in *Lgr5*^DTR-GFP^ organoids from day 0 (seeding) to day 3 in the absence (Control) and presence of 5 mM EIPA (n=3 preparations). Control GFP signal is reduced due to photobleaching. BF, bright-field view. LGR5, *Lgr5*^+^ ISC. (**B**) Quantification of crypt number of day 3 organoids treated without (control) and with 5μM EIPA (n=4; Mann-Whitney test). (**C** and **D**) Crypt budding in NHE1-silenced organoids. (C) Representative images of inducible CRISPR-Cas9 mediated NHE1-silenced organoids from n=7. (D) Quantification of crypt number in control and NHE1-silenced organoids (n=7; Mann-Whitney test). (**E** and **F**) Reversibility of EIPA treatment. (E) Representative images of crypt budding in organoids before and after washing. Pre-wash, 1 day in growth medium (without and with EIPA) after seeding. Post-wash, 1 day after washing and reseeding. (F) Quantification of crypt number in control and EIPA-treated organoids before and after wash (n=3, Mann-Whitney test for each condition; n.s., not statistically significant). (**G**) Representative images of control organoids and organoids lacking NHE1 activity immunolabeled for CD44 to identify crypt cells and lysozyme (LYZ) to identify Paneth cells (n=3). All box plots are minimum to maximum, the box shows 25th-75th percentiles, and the median is indicated as a central line. Data in all box plots with a statistically significant difference are specified with a p value. N.s., not statistically significant. All scale bars represent 20 μm.

### Single-cell RNA-sequencing of intestinal organoids with loss of NHE1 activity

For a global and unbiased view of NHE1 activity-dependent changes in the intestinal epithelium, we performed scRNA-seq on control and NHE1-inhibited small intestinal organoids using the 10x Genomics platform (Methods). After the standard quality control steps (Figure S3A; Methods), we merged data from all 4 conditions (WT+/-EIPA, CRISPR+/-Dox) and performed normalization and batch correction, clustering, cell-type identification, and downstream transcriptional analysis (Luecken and Theis, 2019) (Methods). Each cluster was associated with an enrichment of cell-type specific or cell-cycle signature gene expression, which were then used to specify the identities of each cluster. We observed a strong correlation between our biological replicates (Figure S3B), indicating high reproducibility of our organoid cultures. We visualized the clustering profiles and cell-type gene signatures with UMAP (merged data, Figures 3A and 3B; separated data, Figure S3D) and heatmap (Figure 3C) plots and identified many clusters that were in agreement with previous publications, including generic *Lgr5*^+^ ISCs, enterocytes, tuft cells, enteroendocrine cells (EEC), EEC precursor, Goblet, and Paneth cells (Haber et al., 2017; Qu et al., 2021; Sancho et al., 2015; Tetteh et al., 2016). In addition, we identified two subsets of *Lgr5^+^* ISCs, a low-proliferative and a high-proliferative subpopulations (Figures 3B, 3C and 4A), distinguished by the expression of two reported slowly dividing ISC markers, *Mex3a* (Figure 3C) and *MKi67* (Figure S4A) (Barriga et al., 2017). We also detected differential expression of several other cell cycle genes within these two ISC populations, including *Cdk1*, *Prc1*, and *Cdk6*, and the absorptive lineage-associated gene, *Apoa1* (Figure S4B), consistent with a current view of heterogeneously dividing *Lgr5^+^* ISC pools (Barriga et al., 2017). In the secretory lineage, we identified distinct clusters of progenitors (Castillo-Azofeifa et al., 2019; van Es et al., 2012a; Gregorieff et al., 2009; Lo et al., 2017; Shroyer et al., 2005; VanDussen and Samuelson, 2010), an early secretory progenitor, a precursor of both Paneth and Goblet cells, and a Goblet-specific precursor population (Figures 3B, 3C and 6A; method). We also found a subcluster associated with the enterocyte lineage markers enriched in *Clusterin* (*Clu*) that is involved in the regeneration response in vivo (Figures 3B,3C and 5A; method) (Ayyaz et al., 2019; Lukonin et al., 2020; Qu et al., 2021).

**Figure 3.**
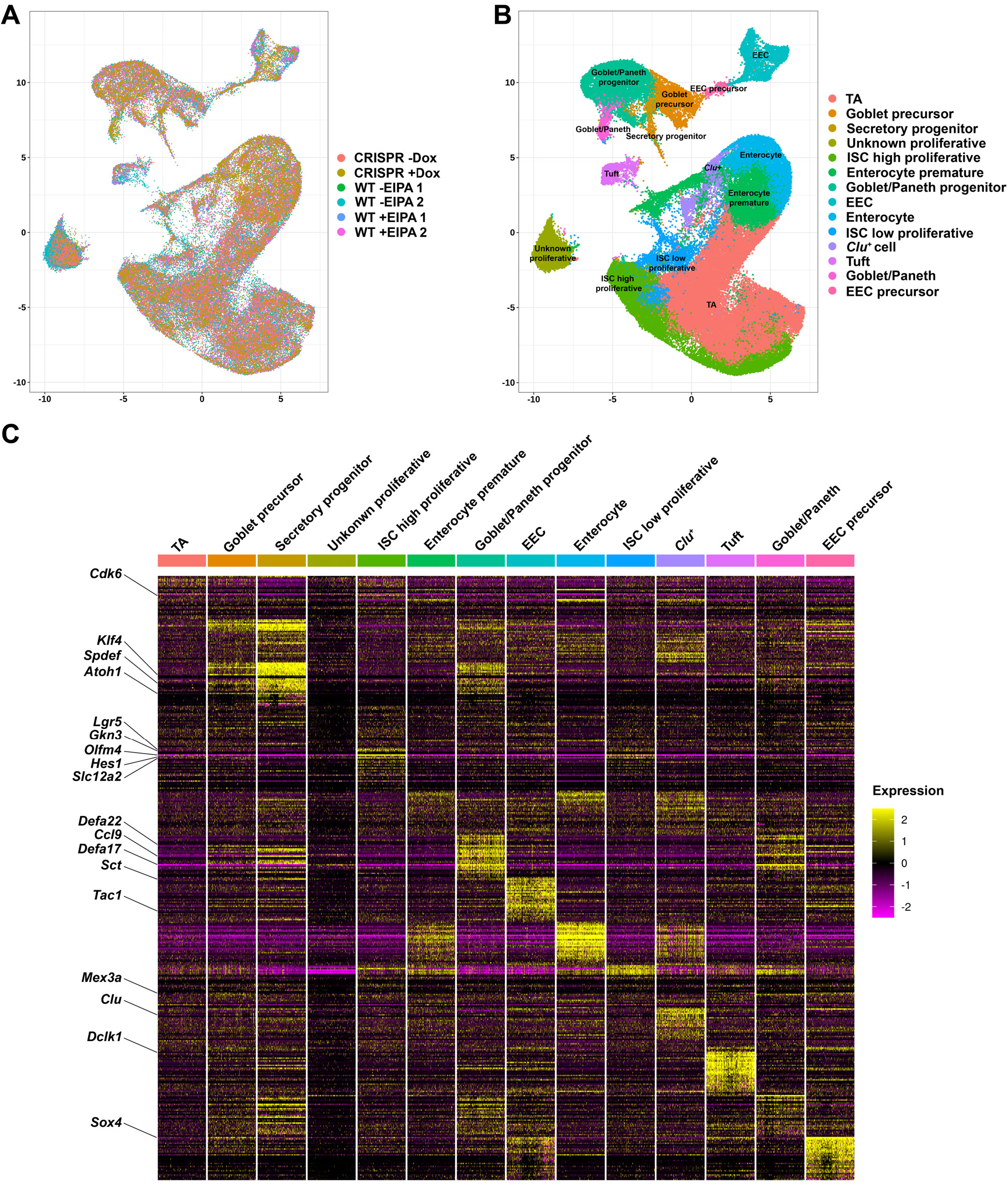
Single-cell transcriptome survey of intestinal organoids. (**A** and **B**) Uniform manifold approximation and projection (UMAP) visualization of cell profiles from a merged dataset (WT+/-EIPA, CRISPR+/-Dox). (A) UMAP of joint single-cell profiles, colored by individual datasets. (B) UMAP of cell-identity clusters based on known markers (Methods), colored by cell identities. (**C**) Heatmap visualization of expression signatures sorted by cell-identity clusters. The colored expression level of each gene per cell, 0 (population mean) ± 2 (Standard Deviation, SD). Conventional cell-type associated genes are highlighted (left row).

### Inhibiting NHE1 reduced crypt base columnar (CBC) cells

ISCs in the crypt base are marked by elevated *Lgr5* expression, and datasets from organoids lacking NHE1 activity contained fewer *Lgr5^+^* ISCs (Figures 4A-4C). In addition, *Lgr5* expression within the ISC cluster was lower than in controls (Figure 4D). Consistent with these data, we found decreased *Lgr5* transcript in EIPA-treated organoids compared with control at day 3 (Figure 4F). We also observed a decrease in GFP expression in *Lgr5-*DTR*-*GFP organoids maintained for 3 days with EIPA compared with controls (Figure 4E; Video S2B). To quantify this effect at the cellular level, we immunolabeled for GFP in *Lgr5*^DTR-GFP^ organoids to identify ISCs and CD44 to identify all crypt cells. We found a significant decrease in the percentage of CD44^+^ cells that are *Lgr5^+^* in organoids maintained with EIPA compared with controls (Figure 4G). The impaired crypt formation in EIPA-treated organoids, however, was not due to a decrease in crypt cell proliferation, because the rate of EdU incorporation in organoids maintained without or with EIPA was not different (Figures 4E and 4H). Together, these observations indicate that inhibiting NHE1 activity causes a decrease in *Lgr5^+^* ISCs without impacting proliferation of crypt cells. Loss of *Lgr5^+^* ISCs is known to cause crypt loss in organoids (Tan et al., 2021) and *in vivo* (van Es et al., 2012b; Fevr et al., 2007; Kuhnert et al., 2004; Pinto et al., 2003). Therefore, the loss of crypt budding with inhibiting NHE1 activity is likely caused by the reduction in *Lgr5^+^* ISCs.

**Figure 4.**
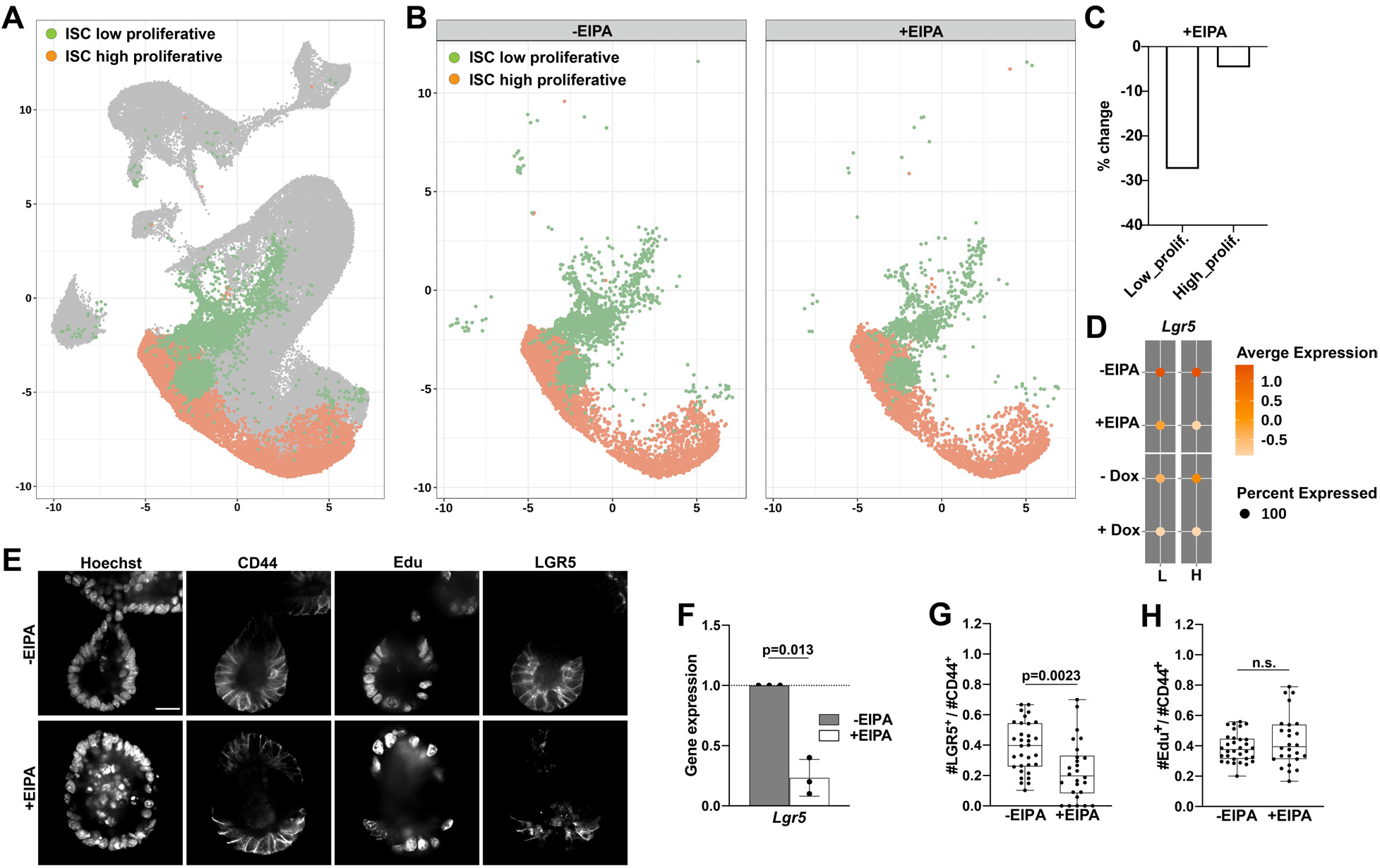
Inhibiting NHE1 activity reduced *Lgr5*^+^ ISCs. (**A**) UMAP plot highlighting distinct *Lgr5*^+^ ISC subtypes. (**B**) Profiles of *Lgr5*^+^ ISC-subtype clusters without (control) and with EIPA. (**C**) Percent change of *Lgr5*^+^ ISC subtypes with NHE1 inhibition by EIPA (Methods). (**D**) Dot plot showing averaged expression level of *Lgr5* in the ISC subtypes. Colored average expression scale, 0 (population mean) ± SD. (**E** to **H**) Crypt proliferation and *Lgr5*^+^ ISC pool with loss of NHE1 activity. (E) Representative images of proliferating (Edu^+^) and *Lgr5*^+^ ISC (LGR5^+^) populations in the crypt region (CD44^+^) of day 3 organoids (n=5). (F) Relative *Lgr5* expression (mean ± SD) in control and EIPA-treated organoids (n=3; Wilcoxon test). Quantification of (G) *Lgr5*^+^ ISCs (LGR5^+^) and (H) number of proliferating cells (Edu^+^) in the crypt region (CD44^+^) (n=5; two-sided student’s t-test; n.s., not statistically significant). All box plots are minimum to maximum, the box shows 25th-75th percentiles, and the median is indicated as a central line. Data in all box plots with a statistically significant difference are specified with a p value. N.s., not statistically significant. All scale bars represent 20 μm.

### Loss of NHE1 activity and decreasing pHi promotes absorptive lineage specification

We next determined whether disrupting the NHE1-generated pHi gradient affects the specification of ISC lineages. The ISC lineage bifurcates into absorptive (enterocyte) and secretory lineages, with the latter including Paneth cells, enteroendocrine cells, and Goblet cells (Beumer and Clevers, 2021; Boonekamp et al., 2020). Comparing the number of cells classified as enterocytes in each scRNA-seq dataset (Figure 5A), we found that the enterocyte cluster constituted a higher percentage of the total cell population in organoids with inhibited NHE1 activity compared with controls (Figures 5B and 5C). Consistent with these data, we observed increased *Alpi1* transcript levels in extracts from organoids with inhibited NHE1 activity compared with controls as determined by qRT-PCR. However, our scRNA-seq data indicated no change in the average expression level of enterocyte signature genes *Alpi1*, *Aldolase B* (*Aldob)*, *Apoa1*, and *Apoa4* in the enterocyte clusters with inhibited NHE1 activity (Figure 6D). This suggests that inhibiting NHE1 activity promotes differentiation toward the absorptive lineage but does not affect the expression of absorptive cell markers in fully differentiated cells. We also observed a significant increase in *Clu* expression within a subset of enterocyte clusters in the datasets from organoids with inhibited NHE1 activity (Figures 5B, 5C, 5E and 5F). Emergence of *Clu*^+^ cells is a signature of intestine regeneration *in vivo* (Ayyaz et al., 2019; Lukonin et al., 2020; Qu et al., 2021), thus raising the possibility that inhibiting NHE1 activity triggers a regeneration-like response in organoids.

**Figure 5.**
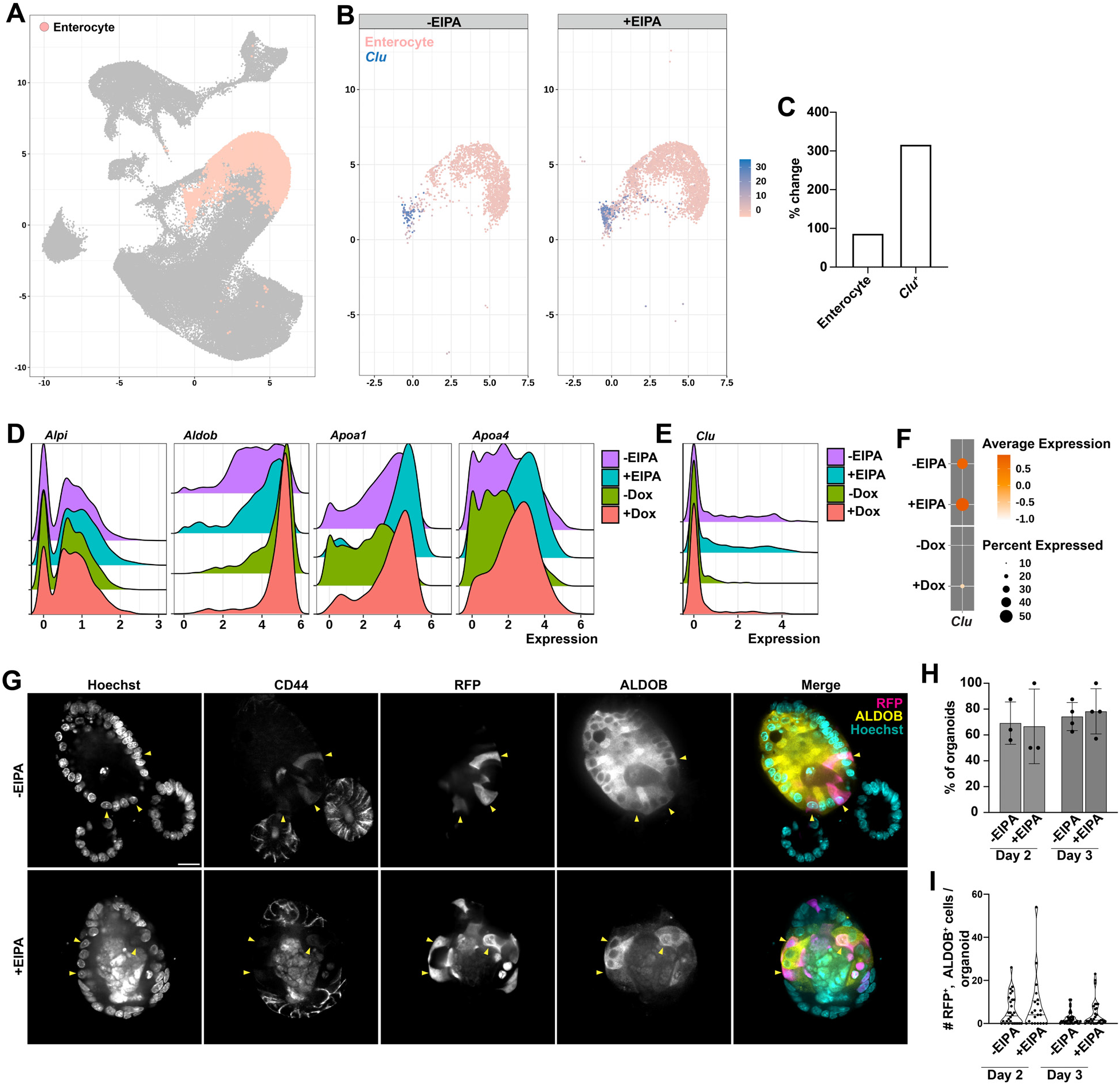
Loss of NHE1 activity promotes the absorptive lineage specification. (**A**) UMAP plot highlighting the enterocyte cluster. (**B**) Profiles of the enterocyte clusters without (control) and with EIPA. The expression of *Clu* (shown as average expression per cell in a scale of log1p) is detected in a subset of the enterocyte clusters. (**C**) Percent change of enterocytes and *Clu*+ cells with NHE1 inhibition by EIPA. (**D-E**) Ridge plots showing the distribution of cells with indicated expression levels of enterocyte markers (D) or *Clu* (E) in the enterocyte clusters with or without NHE1 inhibition. Y-axis, cell counts; X-axis, average expression in a scale of log1p. (**F**) Dot plot showing the *Clu* expression in the enterocyte clusters without and with NHE1 inhibition. Colored average expression, 0 (population mean) ± SD. (**G**-**I**), Lineage tracing for enterocytes in the absorptive cell fate from *Lgr5*^+^ ISCs by using *Lgr5*^CreER^;*Rosa26*^RFP^ organoids. (G) Representative images from 4 preparations show double labeling of *Lgr5*^+^ ISC progeny expressing RFP using and immunolabeled for ALDOB (arrowhead) in the villus region, indicated by CD44^-^, of day 3 organoids without and with EIPA. (H) The percent of organoids (mean ± SD) at which cells in *Lgr5*^+^ ISC labeled villus region (CD44^-^) in day 2 and day 3 organoids are RFP^+^, ALDOB^+^ (day2 n=3, day3 n=4, Mann-Whitney test). (I) Quantification of the number of RFP^+^, ALDOB^+^ double-labeled cells in the villus region (CD44^-^) in day 2 and day 3 organoids maintained in the absence (control) and presence of EIPA (n=4, Mann-Whitney test). All violin plots are minimum to maximum, the dashed line shows 25th-75th percentiles, and the median is indicated as a central line. Data in all violin plots with a statistically significant difference are specified with a p value, otherwise are not significantly different. All scale bars, 20 μm.

**Figure 6.**
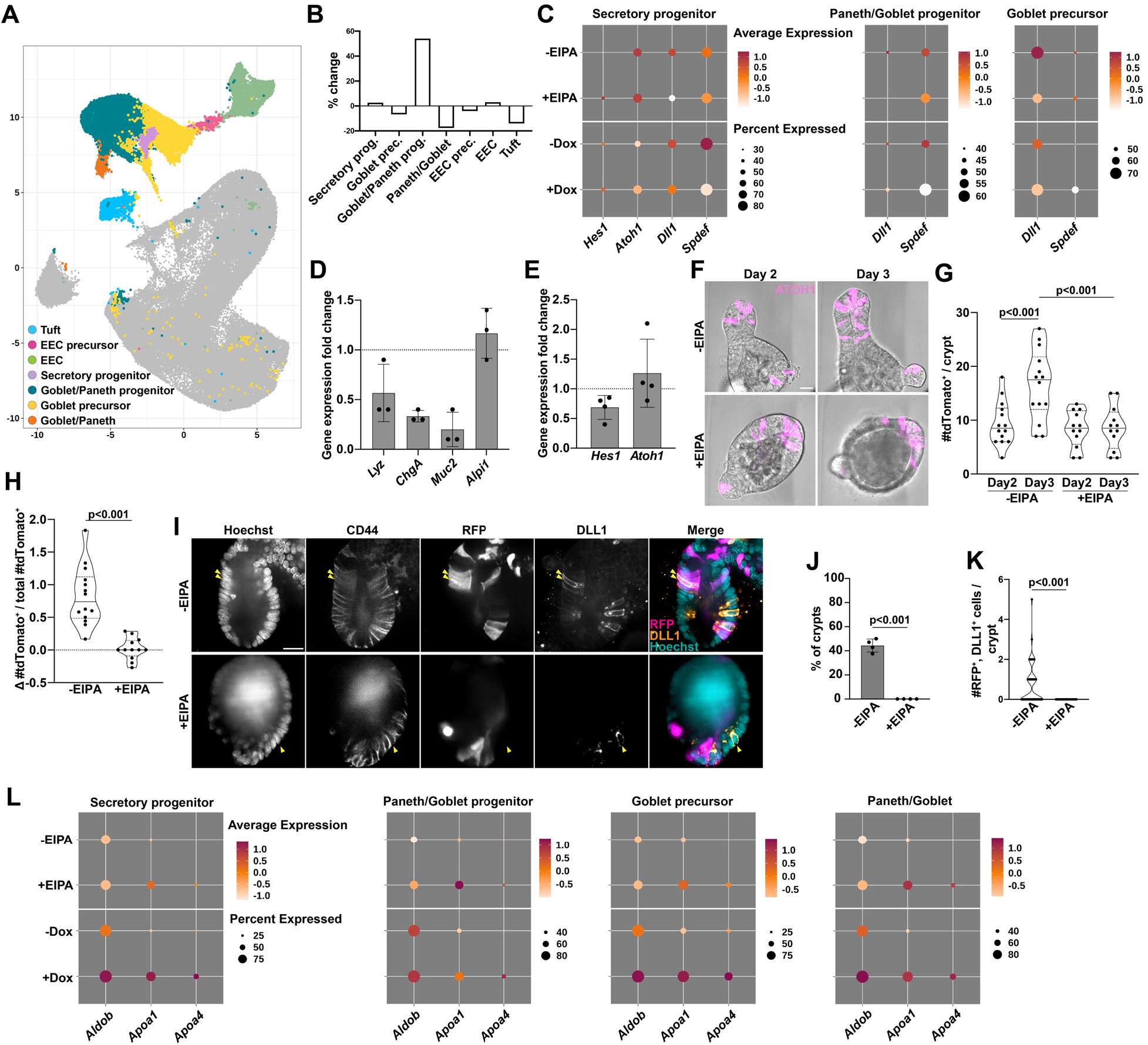
Loss of NHE1 activity impaired the secretory fate by acting on the downstream of ATOH1. (**A**) UMAP plot highlighting secretory clusters. (**B**) Percent changes of secretory cell types with NHE1 inhibition by EIPA. (**C**) Dot plots showing averaged expression level of secretory associated genes in the progenitors and precursor. (**D**) Cell marker expression (mean ± SD) in EIPA-treated organoids relative to controls (n=3). *Lyz*, lysozyme (Paneth cell), *ChgA*, chromogranin A (enteroendocrine cell), *Muc2*, mucin 2 (goblet cell), *Alpi1*, alkaline phosphatase (enterocyte). (**E**) Relative gene expression (mean ± SD) of the Notch signaling pathway in control and EIPA-treated organoids (n=3). (**F** to **H**), Lineage tracing for secretory cells in *Atoh1*^CreERT2^;*Rosa26*^tdTomato^ organoids during development in. the absence (Control) and presence of EIPA. Organoids are untreated or treated with 5 μM EIPA on day 0 (seeding) and then incubated with 4-hydroxytamoxifen added on day 1 followed by 2-day time-lapse imaging to track ATOH1^+^ secretory cells. (F) Representative frames (day 2 and day 3) from a time-lapse recording of 3 preparations show secretory cells indicated by ATOH1^+^ (tdTomato^+^) in organoids without and with EIPA. Time-lapse movie of tdTomato+ secretory cell production (see videos S3A and S3B). (G) Quantification of Atoh1 (tdTomato+) labeled secretory cells from day 2 (first day for lineage tracing) to day 3 within a crypt region in the absence (Control) and presence of EIPA (n=3, two-sided student’s t-test). (H) Changes in the number of newly produced secretory cells in the crypt region in the absence or presence of EIPA. Data are normalized to all labelled secretory cells (tdTomato^+^) seen on day 2 (n=3, two-sided student’s t-test). (**I** to **K**), Lineage tracing for secretory progenitors from *Lgr5*^+^ ISCs using *Lgr5*^CreER^;*Rosa26*^RFP^ organoids. (I) Representative images from 4 preparations, show double labeling of *Lgr5*^+^ ISC progeny expressing RFP and immunolabeled for DLL1 (arrowhead) in the crypt region, indicated by CD44^+^ immunolabeling, of day 2 control but not EIPA-treated organoids. (J) The percent (mean ± SD) of *Lgr5*^+^ ISC labeled crypt region at day 2 organoids containing RFP^+^/DLL1^+^ cells in the absence or presence of EIPA (n=4, Mann-Whitney test). (K) Quantification of the number of RFP^+^/DLL1^+^ cells in the crypt region in day 2 organoids maintained in the absence or presence of EIPA (n=4, Mann-Whitney test). (**L**) Dot plots showing the average expression level of absorptive lineage markers in the secretory cells. All dot plots are colored by average expression, 0 (population mean) ± SD. All violin plots are minimum to maximum, the dashed line shows 25th-75th percentiles, and the median is indicated as a central line. Data in all violin plots with a statistically significant difference are specified with a p value, otherwise are not significantly different. All scale bars, 20 μm.

To further test the effects of EIPA on specification of the absorptive cell lineage, we performed lineage tracing with *Lgr5^CreER^*;*Rosa*26^RFP^ organoids (Castillo-Azofeifa et al., 2019), which traces the differentiation of progeny from *Lgr5^+^* ISCs into all cell types of all major lineages (Figure S5A), including enterocytes (Figure 5G), secretory progenitors (Figure 6I), and Paneth cells (Figure 7A). We observed a slight increase in the number of RFP^+^ enterocytes, marked by *Aldob* expression (Lukonin et al., 2020; Serra et al., 2019) in EIPA-treated organoids at days 2 and 3 compared with controls (Figures 5G-5I). While this difference is not statistically significant, the trend is consistent with the predictions made by scRNA-seq analysis indicating that inhibitng NHE1 activity does not impair differentiation to enterocytes. This observation suggests that there is a robust production of absorptive progenitors in organoids lacking NHE1 activity, which is consistent with our proliferation data (Figure 4E and 4H). Collectively, our data indicate that inhibiting NHE1 activity promotes differentiation toward the absorptive lineage.

**Figure 7.**
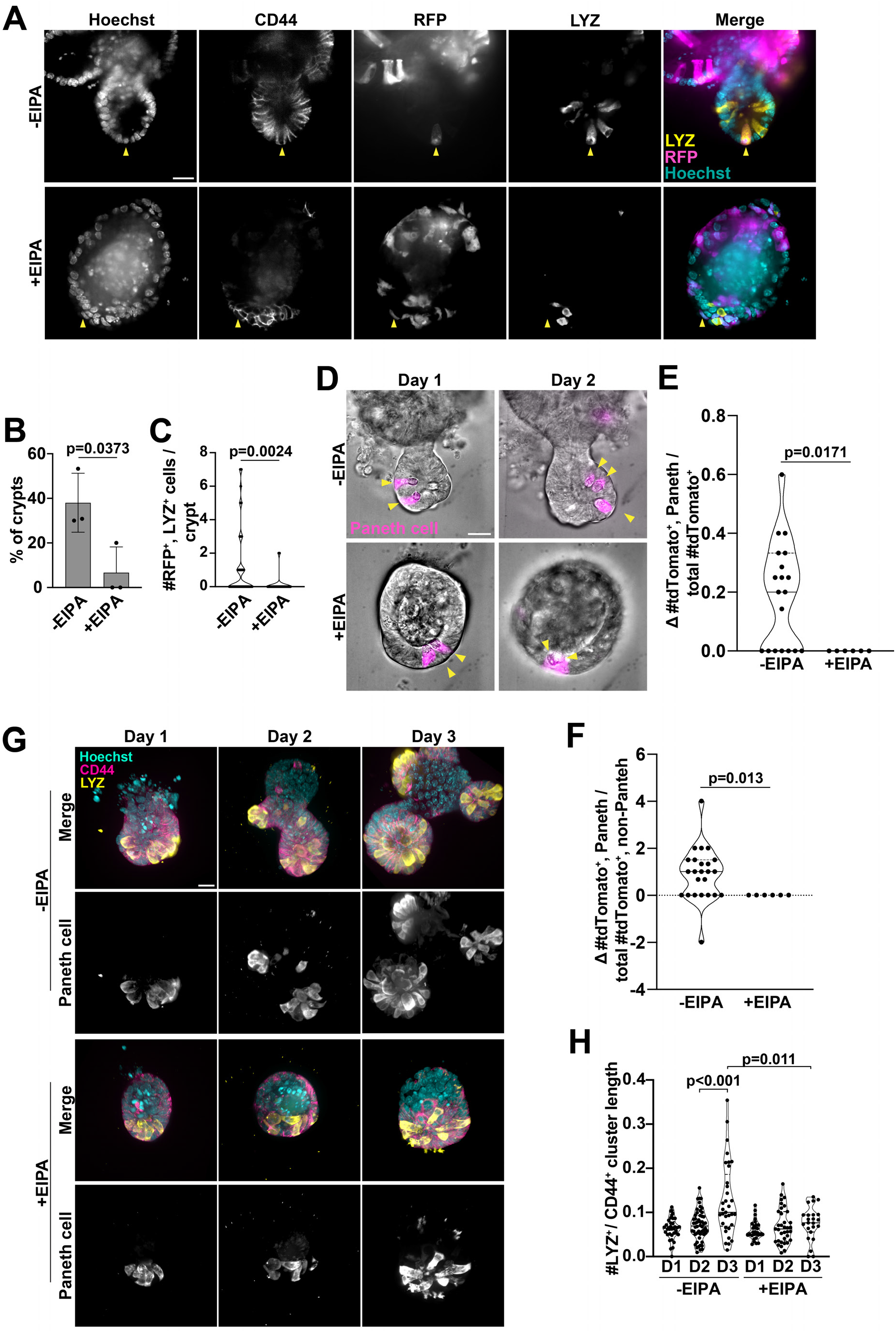
Loss of NHE1 activity impaired Paneth cell differentiation. (**A** to **C**) Lineage tracing for Paneth cells from *Lgr5*^+^ ISCs. (A) Representative images showing the presence of double-labeled *Lgr5*^+^ ISC progeny expressing RFP and immunolabeled with lysozyme (LYZ) antibodies (Paneth cell) (arrowhead) in the crypt region (CD44^+^) of day 3 control but not the EIPA-treated organoids (n=3). (B) Percent of crypt region (mean ± SD) in day 3 organoids contains RFP^+^/LYZ^+^ cells without (control) and with EIPA treatment (n=3, Mann-Whitney test). (C) Quantification of the number of RFP^+^/LYZ^+^ cells in the crypt region of day 3 organoids in the absence and presence of EIPA (n=3, Mann-Whitney test). (**D** to **F**), lineage tracing for newly produced Paneth cells from the secretory progenitors during the development of *Atoh1*^CreERT2^;*Rosa26*^tdTomato^ organoids in the absence and presence of EIPA. Organoids are incubated with 4-hydroxytamoxifen for 24 h, added on day 1 followed by washing and reseeding organoids on day 2. Reseeded organoids are then maintained for 2 days in the absence or presence of 5μM EIPA during which time-lapse imaging is used to track Paneth cell production. (D) Representative frames (day 1 and day 2) from the time-lapse recordings show Paneth cells indicated by the presence of visible dense granules (arrowhead) that are also marked by the secretory lineage marker ATOH1 (tdTomato^+^) to show newly produced Paneth cells from secretory progenitors. (E and F) Changes (from day 1 to day 2) of newly produced Paneth cell number in the crypt region in the absence and presence of EIPA. Data from 3 separate preparations are normalized to either (E) the number of all labelled secretory cells (tdTomato^+^), or (F) the number of non-Paneth secretory cells (tdTomato^+^, non-Paneth) on day 1. Statistical analysis by Mann-Whitney test. (**G** and **H**) Paneth cell abundance in crypts during organoid development from day 1 to day 3. (G) Representative 3D confocal images show Paneth cells in organoid crypts at the indicated days after plating without and with EIPA (n=3). CD44, crypt region. LYZ, lysozyme, Paneth cell. (H) Quantification of Paneth cell numbers from conditions described (G) (n=3; Mann-Whitney test and two-sided student’s t-test). D, day. All violin plots are minimum to maximum, the box shows 25th-75th percentiles, and the median is indicated as a central line. Data in all box plots with a statistically significant difference specified with p values, otherwise are not significantly different. All scale bars represent 20 μm.

### Lowering pHi impairs the secretory fate by acting downstream of ATOH1

We used a combination of scRNA-seq and lineage-tracing to determine whether the loss of NHE1 activity affects specification of the secretory lineage. The scRNA-seq datasets indicated that organoids lacking NHE1 activity had a higher percentage of cells classified as secretory progenitors or Paneth-Goblet progenitors and a lower percentage of differentiated Paneth-Goblet and Tuft cells compared with controls (Figures 6A and 6B), suggesting impaired specification of secretory cells. Because specification of the secretory lineage is regulated by Notch signaling pathway activity in the daughter cells of ISCs (Fre et al., 2005; Sancho et al., 2015; Stanger et al., 2005; VanDussen and Samuelson, 2010; VanDussen et al., 2012), we investigated whether inhibiting NHE1 activity influences Notch pathway in the ISC lineage. It is well-documented that Notch lateral inhibition causes immediate daughter cells from ISCs to adopt a state of either high or low Notch pathway activity. High levels of Notch signaling promote the expression of *Hes1*, which represses the master secretory lineage transcription factor ATOH1 and thus promotes differentiation toward the absorptive cell fate. Conversely, in cells with low Notch pathway activity, low levels of HES1 permits *Atoh1* expression, which promotes the secretory cell fate. ATOH1 forms a positive feedback loop by reinforcing its own expression and promoting the expression of *Dll1* and *Dll4*, thus enabling lateral inhibition of the secretory fate in neighboring cells (van Es et al., 2005; Fre et al., 2005; Sancho et al., 2015; Stanger et al., 2005; Tomic et al., 2018; VanDussen and Samuelson, 2010; VanDussen et al., 2012). *Dll1* and *Spdef*, are ATOH1 targets that are known to be involved in the secretory cell differentiation (Gerbe et al., 2011; Gregorieff et al., 2009; Lo et al., 2017; Noah et al., 2010; Shroyer et al., 2005). We found that expression of *Hes1* and *Atoh1* was similar in secretory progenitor cell populations from control and experimental datasets (Figure 6C), suggesting the Notch pathway activity was not affected by inhibiting NHE1 activity. However, the targets of ATOH1, including *Dll1* and *Spdef*, were generally decreased (Figure 6C). Likewise, we also observed decreased expression of these genes in both the Paneth-Goblet progenitor and the Goblet precursor populations (Figure 6C), supporting our observation that the secretory cell population but not the ATOH1^+^ secretory progenitor cell population is reduced (Figure 6B). We also found that several genes typically associated with the absorptive fate were elevated in multiple secretory cell types with loss of NHE1 activity, including the secretory progenitors and the Goblet cell precursors (Figure 6L). These findings suggest that NHE1 activity helps to reinforce the distinction between the absorptive and secretory lineages, and that inhibiting NHE1 activity impairs secretory lineage specification.

To further test the prediction that lowering pHi impairs downstream function of ATOH1 and secretory lineage specification, we profiled global changes of secretory markers by qRT-PCR and found that expression of all three secretory cell markers – *Lyz* for Paneth cells, *ChgA* for enteroendocrine cells, and *Muc2* for Goblet cells – were decreased with loss of NHE1 activity compared with control (Figure 6D). In addition, we observed no statistically significant change in *Hes1* and *Atoh1* expression in EIPA-treated organoids compared with controls (Figure 6E). These data suggest that the block in secretory cell fate specification with loss of NHE1 activity is due to an effect that is downstream of Atoh1 expression. To test this hypothesis, we performed lineage tracing with *Atoh1*^CreERT2^; *Rosa26*^tdTomato^ organoids (Castillo-Azofeifa et al., 2019) (Figure 6F; Videos S3A and S3B). We found no differences in the number of tdTomato^+^ cells in control and the EIPA-treated organoids on Day 1 of lineage tracing (Figure 6G), indicating the induction of clones from ATOH1^+^ cells was similar in both conditions. However, whereas there was a net increase of tdTomato^+^ cells over time in control organoids, the number of tdTomato^+^ cells per crypt remained constant in organoids treated with EIPA (Figures 6G and 6H; Videos S3A and S3B). As an additional test of the effect of NHE1 inhibition on secretory cell fate specification, we performed lineage-tracing with *Lgr5^CreER^*;*Rosa*26^RFP^ organoids and assayed for RFP^+^ cells that had differentiated into DLL1^+^ secretory progenitors (Figure 6I). We found that the frequency (Figure 6J) and number (Figure 6K) of RFP^+^, DLL1^+^ cells in the CD44^+^ region were significantly reduced in EIPA-treated organoids compared with controls. Taken together, these data indicate that loss of NHE1 activity impairs differentiation of ATOH1^+^ secretory progenitor into mature secretory cells.

### Loss of NHE1 activity impairs Paneth cell differentiation

Paneth cells are a major cell type in the secretory lineage and are required for ISC self-renewal in organoid cultures. Therefore, we tested the prediction that loss of NHE1 activity impairs Paneth cell differentiation by generating clones in *Lgr5^CreER^*;*Rosa*26^RFP^ organoids and immunolabeling for the Paneth cell marker, LYZ. Indeed, we found loss of NHE1 activity significantly decreased the number of LYZ^+^ Paneth cells in the RFP^+^ clones generated from *Lgr5^+^* ISCs (Figures 7A-7C). Even in EIPA-treated organoids in which nearly all the cells were RFP^+^, Paneth cells remained negative for RFP (Figure S5B). These findings illustrate that, in the presence of EIPA, Paneth cell production by *Lgr5^+^* ISC is impaired. To further test how inhibiting NHE1 affects Paneth cell fate from the ATOH1^+^ secretory progenitor, we quantified Paneth cell production by tracking the *Atoh1*^CreERT2^; *Rosa26*^tdTomato^ organoids from day 1 to day 2 and traced the tdTomato^+^ cells containing dense granules, which are visible with bright-field microscopy (Figure 7D). In control organoids, the number of Paneth cells significantly increased from day 1 to day 2, as expected, whereas with EIPA the number of tdTomato^+^ Paneth cells remained constant (Figures 7D-7F). Hence, inhibiting NHE1 activity impaired Paneth cell differentiation from an ATOH1^+^ secretory progenitor.

As an additional test, we immunolabeled for CD44 and the Paneth cell marker, LYZ (Figure 7G) using organoids maintained without or with EIPA. The number of LYZ^+^ cells significantly increased from day 1 to day 3 in control organoids but remained unchanged over this period with EIPA (Figure 7H). This demonstrates that new Paneth cells accumulate in the growing crypt under normal conditions, consistent with the idea that constant replenishment is important to maintain the Paneth cell population in the crypt, and that NHE1 activity is required for this process. Taken together, our findings indicate that loss of NHE1 activity impairs Paneth cell fate specification, therefore decreasing the Paneth cell replenishment and reducing Paneth cell numbers in the growing crypt.

### Exogenous WNT rescues crypt budding impaired with loss of NHE1 activity

In organoids maintained under standard growth conditions without WNT in the medium Paneth cells are the only source of WNT ligands for ISCs (Gehart and Clevers, 2019; Sato et al., 2011). Thus, to test whether loss of NHE1 activity impaired WNT signaling in organoids, we first immunolabeled for EPHB2, which is activated by WNT signaling and expressed throughout the crypt (Batlle et al., 2002; Holmberg et al., 2006). Indeed, as predicted by our observation that inhibition of NHE1 activity reduces Paneth cell number, we observed a substantial decrease in EPHB2 labeling in EIPA-treated organoids compared to controls (Figure 8A; Figure S6B). However, EIPA-treated organoids are not completely devoid of Paneth cells so, to test whether this effect is also due to an inability of the existing Paneth cells to induce WNT signaling in ISCs, we performed a single cell-based *in vitro* reconstitution assay (Rodríguez-Colman et al., 2017; Sato et al., 2011; Yilmaz et al., 2012). We found that mixing single WT *Lgr5^+^* ISCs with single NHE1-silenced Paneth cells in the absence of exogenous WNT3A did not reduce the efficiency of organoid formation (Figure S6A). This suggests that the reduction in WNT signaling is due to the decreased Paneth cell number rather than an impaired ability of mature Paneth cells to secrete WNT ligands.

**Figure 8.**
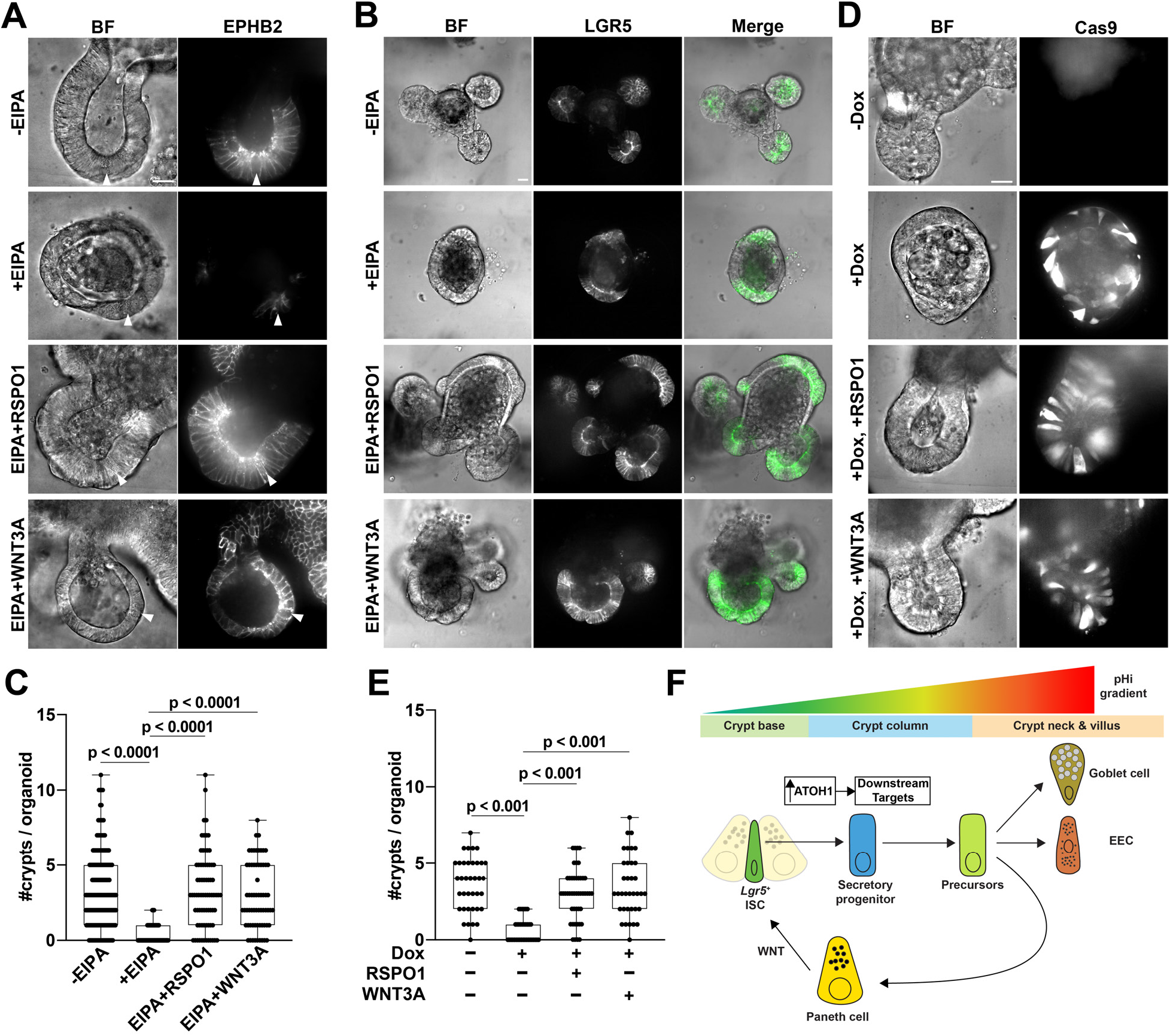
Exogenous WNT rescues crypt budding impaired with loss of NHE1 activity. (**A**) Exogenous WNT restores EPHB2 impaired by NHE1 inhibition. Representative confocal images from 3 cell preparations of EPHB2 immunolabeled organoids in the absence of EIPA (control), the presence of EIPA, or with EIPA plus increased Rspondin1 or exogenous WNT3A. BF, brightfield; arrowheads, Paneth cells in a crypt region. Quantified EPHB2 immunolabeling (see Figure S6B). (**B** to **E**) Representative confocal images and quantification of budded crypts at day 3 in *Lgr5*^DTR-GFP^ organoids with NHE1 inhibition in the presence of exogenous WNT. (B and C) Crypt growth in (B and C) the WT organoids in the absence (control) and presence of EIPA, and in (D and E) the inducible CRISPR-Cas9 organoids without (control) and with Dox, and supplemented with increased RSPO1 or WNT3A. BF, bright-field view; LGR5, *Lgr5*^+^ ISC. Data are from 5 preparations in the WT organoids, and 3 preparations with the inducible CRISPR-Cas9 organoids. Data are statistically analyzed by the Mann-Whitney test. (**F**) Schematic representation of the main findings. All box plots are minimum to maximum, the dashed line shows 25th-75th percentiles, and the median is indicated as a central line. Data in all box plots with a statistically significant difference are specified with a p value, otherwise are not significantly different. All scale bars, 20 μm.

We previously reported that decreased pHi cell-autonomously enhances WNT pathway activity by stabilizing β-catenin (White et al., 2018). Therefore, although WNT signaling is decreased overall in organoids with a loss of NHE1 activity due to the reduction in Paneth cell number, we considered the possibility that loss of NHE1 activity may cause a cell-autonomous increase in WNT signaling within the ISCs. Indeed, we found that expression of canonical WNT pathway targets, *Axin2* and *Ascl2*, was slightly increased in the ISCs from organoids with inhibited NHE1 (Figure S7A). Likewise, we observed a similar increase of these canonical WNT targets by qRT-PCR in organoids treated with EIPA, but no difference in the non-WNT responsive ISC marker *Olfm4* (van der Flier et al., 2009; VanDussen et al., 2012) (Figure S7B) compared with controls. Thus, loss of NHE1 activity does not interfere with the ability of ISCs to induce WNT target gene expression and, in fact, may induce a further increase in the expression of these genes. Taken together with our finding that inhibiting NHE1 does not impair the ability of Paneth cells to promote ISC self-renewal (Figure S6A), these results suggest that the decrease in ISC number and impaired crypt budding in the absence of NHE1 activity is likely a direct consequence of the loss of Paneth cell differentiation.

As Paneth cells provide essential WNT ligands to ISCs, we next tested whether restoring WNT signaling is sufficient to rescue the ISC maintenance and crypt budding phenotypes caused by inhibition of NHE1 activity. We found that ISC number was substantially increased and crypt budding was restored by the addition of either 20% WNT3A ligand or RSPO1 to the culture medium of these organoids (Fig. 8A-E). Notably, 20% WNT3A is a relatively low concentration that is not sufficient to induce spheroids. In addition, we observed that production of the WNT-responsive gene EPHB2 was restored to control levels, which confirms that WNT signaling is abrogated by inhibiting NHE1 activity (Figure 8A; Figure S6B). Taken together, our findings support a model in which an NHE1 activity-dependent pHi gradient that acts downstream of ATOH1 function, facilitates crypt homeostasis by regulating lineage specification (Figure 8F).

## Discussion

We identified a pHi gradient in mouse small intestinal organoids, with a lower pHi in cells at the crypt base and a progressive increase along the crypt column. Attenuating this gradient by pharmacologically or genetically inhibiting activity of NHE1, a plasma membrane H^+^ extruder, inhibited the cell-fate decision of *Lgr5*^+^ ISCs toward the secretory lineage and biased it toward the absorptive lineage. This altered lineage specification occurs in part by reducing ATOH1-dependent signaling and thus decreasing Paneth cell differentiation, which is essential for crypt budding. Moreover, these findings were indicated by scRNA-seq datasets and confirmed experimentally by immunolabeling, single cell reconstitution, and lineage tracing.

Increasing evidence indicates a previously unrecognized role for pHi dynamics in the differentiation of diverse types of stem cells. Increased pHi is necessary for the differentiation of clonal mouse embryonic stem cells from naïve to primed states (Ulmschneider et al., 2016), mesenchymal stems cells to cardiomyocytes (Li et al., 2009), and CD4^+^ T helper 9 cells (Singh et al., 2016). Additionally, *in vivo* studies confirm that increased pHi is necessary for differentiation of *Drosophila* follicle stem cells (Benitez et al., 2019; Ulmschneider et al., 2016), melanocytes during zebrafish neural crest development (Raja et al., 2020), and mesoderm progenitors in the chicken embryo (Oginuma et al., 2020). Similar to our findings, an increasing pHi gradient is seen in murine colonic crypts (Amiri et al., 2021; Nikolovska et al., 2022), and reducing pHi is associated with decreased expression of absorptive fate genes (Nikolovska et al., 2022). Our study is distinct from these reports and adds to this growing area of investigation by identifying a role for pHi dynamics in lineage specification and providing a mechanistic understanding of pHi dynamics regulating ATOH1-dependent fate decision.

Our findings are consistent with roles for pH in regulating WNT signaling and provide new insight into the crosstalk between pHi dynamics and cell signaling. Specifically, decreased vacuolar pH promotes canonical WNT signaling by increasing LRP activation (Cruciat et al., 2010), and deceased cytoplasmic pH (pHi) regulating Dishevelled localization (Simons et al., 2009), and β-catenin acetylation (Oginuma et al., 2020) and stability (White et al., 2018). Thus, the pHi gradient in intestinal crypts may help shape the pattern of WNT signaling, which is highest at the crypt base where pHi is low and decreases along the crypt axis where pHi is higher (Farin et al., 2016; Kosinski et al., 2007; van de Wetering et al., 2002). In addition, our study provides evidence that pHi dynamics regulate cell-fate decisions downstream of a separate signaling cue, ATOH1, which is a master regulator of the secretory lineage. This likely occurs within the progenitor cells that first upregulate ATOH1 to specify differentiation toward the secretory rather than absorptive fate. In addition, our observation that genes typically associated with the absorptive cell fate were upregulated in multiple *Atoh1*^+^secretory cell types when pHi gradient is disrupted suggests that pHi dynamics continues to stabilize the secretory cell fate choice even after the initial lineage specification. Signaling by pHi dynamics is often mediated by intrinsic pH-sensing proteins with activity or protein-protein and protein-phospholipid binding affinities regulated within the cellular pH range (Schönichen et al., 2013). In line with this view, ATOH1 may be a pH sensor with increased pHi enabling its transcriptional function. Alternatively, other pHi-sensitive proteins may be acting to modify ATOH1 activity.

Our findings with disrupting the pHi gradient are consistent with recent studies of plasticity of small intestinal epithelium, further demonstrating the flexibility of pHi as regulator for cell behaviors. Specifically, our finding that with loss of NHE1 activity there is an increase in the number of *Clu*^+^ cells, which is a signature of intestinal regeneration, suggests that inhibiting NHE1 activity or decreasing pHi may elicit a response that mimics tissue regeneration induced by intestine injury *in vivo* (Ayyaz et al., 2019; Lukonin et al., 2020; Qu et al., 2021; Sprangers et al., 2021). Hence, targeting NHE1 activity may be an alternative approach to current methods such as the use of radiation for investigating the acute tissue regeneration in the intestinal epithelium. In addition, our studies are consistent with current views about how crypt budding is regulated. Specifically, current views suggest that crypt budding is maintained by a constant pool of ISCs (van Es et al., 2012b; Tan et al., 2021), which is in line with our findings here, or by actin remodeling that drives a change in mechanical forces (Beuvery et al., 1986; Sumigray et al., 2018; Yang et al., 2021), which has been shown to be regulated by pHi in other contexts (Donahue et al., 2021; Frantz et al., 2008; Webb et al., 2016).

Our findings also have clinical relevance. Cystic fibrosis (CF) is associated with increased intestinal epithelial cell proliferation and an increased risk for gastrointestinal tumors. Increased pHi is seen with CF, and recent reports indicate that impaired intestinal homeostasis is in part mediated by disrupted pHi dynamics (Liu et al., 2012; Strubberg et al., 2018), including a dysregulated pool of ISCs and Paneth cells as we found. Additionally, constitutively increased pHi is seen in most cancers (Webb et al. 2011; White et al. 2017), which could promote expansion of *Lgr5*^+^ stem cells critical for initiating gastrointestinal cancers (Barker et al., 2009; Fatehullah et al., 2021; Shimokawa et al., 2017). Consistent with this possibility, decreasing pHi by genetic silencing of NHE1 or carbonic anhydrase 9 significantly reduces the growth of colon cancer cells (Parks et al., 2017). Additionally, most colorectal cancers are caused by mutations in key components of the WNT pathway, which would avert pHi regulated WNT pathway activity. Altogether, our finding of pHi regulating ATOH1-dependent ISC lineage specification that connects ISC maintenance and the WNT circuit (Figure 8F) adds new insight into the development of CF and intestinal tumorigenesis and provides a biological basis for targeting NHE1 and pHi dynamics for potential therapeutics.

## Funding

National Institutes of Health grant CA197855

National Science Foundation grant P0538109

National Science Foundation grant 1933240

UCSF Research Allocation Program

UCSF RAP Pilot for Established Investigators in Basic and Clinical/Translational Sciences

National Institutes of Health, Maximizing Investigators’ Research Award DE026602

## Author contributions

Conceptualization: Y.L., T.N., D.L.B.

Experiment design and data analysis: Y.L., T.N., D.L.B., E.R., D.C., O.D.K.

Investigation: Y.L.

Funding acquisition: T.N., D.L.B., O.D.K.

Supervision: T.N., D.L.B.

Writing – original draft: Y.L., T.N., D.L.B.

Writing – review & editing: Y.L., T.N., D.L.B., E.R., D.C., O.D.K.

## Data and materials availability

The scRNA-seq data can be retrieved from NCBI gene expression omnibus (GEO). Codes for bioinformatic analysis is deposited in Github repository. All other data are available in the main text and the supplementary materials.

## Methods

### DNA constructs and CRISPR design

The pLX304 lenti-mCherry-SEpHluorin plasmid was generated by inserting an mCherry-SEpHluorin fragment from an mCherry-SEpHluorin plasmid (Addgene #32001) into lentiviral vector pLX304 (Addgene #25890). The mCherry-SEpHluorin fragment was first amplified and then cloned into the Gateway entry vector pENTR/D-TOPO (Thermofisher) to build pENTR/D-mCherry-SEpHluorin. The mCherry-SEpHluorin fragment was then transferred from pENTR/D-mCherry-SEpHluorin into pLX304 via LR gateway reaction to build pLX304 lenti-mCherry-SEpHluorin. To generate the pTLCV2 NHE1 KO plasmid, guide RNA targeting the first exon of the mouse NHE1 gene was designed and cloned into an all-in-one doxycycline (Dox)-inducible vector TLCV2 (Addgene #87360). In brief, gRNA oligo forward (5’-caccgAACTTAATCATTGAACATGG-3’) and reverse (5’-aaacCCATGTTCAATGATTAAGTTc-3’) were first annealed and then ligated into BsmBI digested TLCV2 vector.

### Small intestinal organoids and organoid culture

WT, *Lgr5*^DTR-GFP^ organoids (Tian et al., 2011), *Lgr5*^CreER^;*Rosa*26^RFP^ organoids (Castillo-Azofeifa et al., 2019), and *Atoh1*^CreERT2^;*Rosa26*^tdTomato^ organoids (Castillo-Azofeifa et al., 2019) were obtained from adult mice. The mCherry-SEpHluorin and inducible NHE1 CRISPR-Cas9 KO organoids were generated by infecting WT organoids with lentivirus made from pLX304 lenti-mCherry-SEpHluorin and pTLCV2 NHE1 KO plasmids, respectively, followed by antibiotic selection using 10 μg/mL blasticidin for mCherry-SEpHluorin and 2 μg/mL puromycin for NHE1 CRISPR-Cas9 organoids, as described (Koo et al., 2011). Single clones of NHE1 silenced organoids were established from selected individual puromycin-resistant organoids. The organoid cultures were generated and maintained as previously described (Koo et al., 2011; Mahe et al., 2013; Sato et al., 2009). Briefly, organoids were embedded in Matrigel (Corning, 356231) and cultured in a standard 24-well plate. For confocal imaging, organoids embedded in Matrigel were seeded into 24-well glass-bottom MatTek plates (MatTek, P24G-0-10-F). Unless stated otherwise, organoids were grown in ENR medium containing advanced DMEM/F12 (Invitrogen, 12634-028) supplemented with 50 ng/mL EGF (Sigma-Aldrich, E9644-.2MG), 100 ng/mL Noggin (R&D, 6057-NG/CF), R-spondin1(RSPO1) (conditioned medium, CM; CCHMC RT0457, 1.6% v/v), 10 mM HEPES (Invitrogen, 15630-080), 1 mM N-acetylcysteine (Sigma-Aldrich, A7250), 1X glutaMAX (Invitrogen, 35050-061), 1X N2 supplement (Invitrogen, 17502-048), 1X B27 supplement (Invitrogen, 17504-044), and 1X penicillin-streptomycin (pen/strep). To induce spheroid formation, organoids were cultured in ENRWN medium, which includes ENR medium with 50% v/v WNT3A (CM, ATCC CRL-2647) and 10 mM Nicotinamide (Sigma-Aldrich, N1630-100MG). For post-infection cultures, medium with an additional 10 μM of the Rho kinase inhibitor Y-27632 (Sigma-Aldrich, Y0503-1MG) in ENRWN was used for recovery. For WNT rescue, ENR medium was supplemented with 6.4% v/v RSPO1 (CM, CCHMC RT0457) or 20% v/v WNT3A (CM, ATCC CRL-2647). For single-cell reassociation assays, ISC-Paneth pairs were grown in single-cell growth medium, which was ENR with 10 mM Nicotinamide, 2.5 μM Y-27632, 2.5 μM Chir99021 (Sigma-Aldrich, SML1046-5MG), 1 μM Jagged-1 (Anaspec, AS-61298), 2.5 μM Thiazovivin (Selleckchem, S1459).

### NHE1 CRISPR-Cas9 silencing and validation

To induce and validate NHE1 CRISPR-Cas9 editing, 2 μg/mL doxycycline (Dox) was added to the ENR medium and to Matrigel after passaging organoids. After induction for 2-3 days, clones containing at least 80% Cas9-EGFP positive organoids were selected for genomic DNA (gDNA) extraction. The gDNA was isolated from selected clonal organoids using the gDNA tissue miniprep system (Promega, A2051) and evaluated by PCR and DNA gel electrophoresis using primer pair forward (5’-GCCCGTGGTCCAGCCTATC-3’) and reverse (5’-GTCCCATCCCAGCTGTAGGAGA-3’). Purified PCR fragment containing the edited DNA was sequenced by Sanger sequencing using primer forward (5’-CGTCTGGGGATTTCATCCACCT-3’) and reverse (5’-CTATCTTCATGAGGCAGGCCAGGA-3’) respectively. The sequencing results were analyzed by comparing WT and -Dox samples (Figure S2).

### pHi determination

Ratiometric imaging of the pHi biosensor mCherry-SEpHluorin was performed as described (Grillo-Hill et al., 2014). In brief, organoids embedded in Matrigel were cultured in ENR medium supplemented with DMSO vehicle (1:1,000) or 5 μM EIPA for 1 to 3 days. Before imaging, growth medium was replaced with freshly made pHi buffer containing 25 mM NaHCO_3_, 115 mM NaCl, 5 mM KCl, 10 mM glucose, 1 mM K_3_PO_4_, 1 mM MgSO_4_, and 2 mM CaCl_2_ pH 7.4 and including the membrane dye CellMask™ (Molecular probes, C10046) for 10 min. After washing with fresh pHi buffer not containing CellMask™ fluorescence images were acquired using a customized spinning disk confocal (Yokogawa CSU-X1) on a Nikon Ti-E microscope equipped with a live-cell imaging chamber maintained with 5% CO_2_ at 37°C, a 40X water objective, 488 nm, 560 nm, and 590 nm excitation lasers. and a Photometrics cMYO cooled CCD camera. Images were stored as Z-stacks of 2 μm-optical sections. To calibrate fluorescence ratios of mCherry-SEpHluorin to pHi, after each experiment, organoids were incubated for 15-20 min with a KCl buffer (80 mM KCl, 50 mM K_3_PO_4_, 1 mM MgCl_2_) containing 20 μM nigericin (Invitrogen, N1495) at pH 7.8. After acquiring fluorescence ratios, organoids were washed and incubated with nigericin buffer at pH 6.6 and fluorescence ratios were again acquired to generate a 2-point calibration conversion.

For image analysis, backgrounds were removed from Z stacks and single optical sections containing representative intestinal crypts were chosen for quantification. Individual cells were segmented by the membrane dye. To identify major cell types in the crypt, Paneth cells were selected based on size and presence of visible intracellular granules under both bright-field and mCherry channels, ISCs were identified based on their location which is intermingled between two Paneth cells, other cells along the crypt column starting at a position two-cell above the upmost Paneth cell and ending at the crypt neck, the upmost part of a budded crypt, were further specified as the crypt column and the crypt neck. In the crypt without a bud, up to four of the cells closest to the upmost Paneth cell were chosen as the column for pHi measurement. Because the crypt region and villus region were less distinguishable by live-cell imaging in the day 1 organoid during development, the column region was not specified for pHi determination on day 1. To calculate mCherry/SEpHluorin intensity ratios, fluorescence intensity was measured in the selected individual Paneth cells, ISCs, and cells located in the column and neck using ImageJ. In Microsoft Excel, the pHi values were determined by applying those intensity ratios to a standard curve generated by graphing the nigericin pH values (Y-axis) against their dependent intensity ratios (X-axis). Because of the mCherry/SEpHluorin intensity ratio in the cells in the organoid villus (above the crypt neck) was too high to be calibrated, the pHi in those cells were not determined.

### Crypt budding quantification and imaging

Under a bright-field view on the spinning disc confocal microscope, a budded crypt was defined by the epithelial protrusion (containing visible Paneth cells) grown out of the spherical cyst with the extension protruded larger than the thickness of the epithelial layer of the cyst. To quantify crypt budding, the number of budded crypts were counted in organoids under each of the following conditions. (1) For -EIPA vs +EIPA conditions, day 3 organoids grown in ENR medium supplemented with DMSO (vehicle control) or 5 μM EIPA were scored for crypt budding. (2) For inducible CRISPR-Cas9 mediated NHE1 silencing, organoids in ENR medium were treated with or without Dox for 2 days and then re-seeded in ENR with or without Dox for another 3 days before quantification. (3) For WNT rescue experiments, EIPA-treated and CRISPR-Cas9 silenced organoids were cultured in the ENR medium supplemented with either 3X RSPO1 or 20% exogenous WNT3A for 3 days prior to quantification. To imaging crypt budding in the *Lgr5*^DTR-GFP^ organoids (Figure 2A), a 3-day time-lapse imaging (bright-field, FITC; 40X) began immediately after seeding (day 0) in ENR + DMSO (vehicle control) and ENR + 5 μM EIPA. Images were acquired as Z-stacks of 2 μm optical sections every 3 hrs. To image pHi dynamics in crypt budding in the mCherry-SEpHluorin organoids, a time course was acquired (bright-field, FITC, PE-Texas red; 20X) was starting one day after seeding, in which a 2 μm Z-stack image set was taken every 30 mins for two days.

### Single-cell RNA-sequencing (scRNA-seq) of intestinal organoids

To obtain single-cell suspension from the organoids without or with NHE1 inhibition (-/+ EIPA, -/+ Dox), organoid cultures were first washed with the cold (4°C) advanced DMEM/F12 (Invitrogen, 12634-028) supplemented with 10 mM HEPES (Invitrogen, 15630-080), 1 mM N-acetylcysteine (Sigma-Aldrich, A7250), 1X glutaMAX (Invitrogen, 35050-061), and 1X pen/strep. After the washing, wells containing the organoids were incubated with 300 uL cold (4°C) Corning recovery solution (Corning, 354253) for 15 min to melt the Matrigel. Next, organoids were mechanically disrupted with the P1000 pipette tips first and then trypsinized with RT TrypLE Express (Gibco, 1952062) for up to 15 min to disassociate into single cells. Single cells were then suspended within a sorting buffer that contained Hanks’ balanced salt solution (HBSS) with 3% m/v FBS, 10 mM HEPES and 5 mM EDTA. After staining with Live/Dead (Invitrogen, L23105, 1:1000) in the sorting buffer, fluorescence-activated cell sorting (FACS) was performed on the BD FACSAira instrument to enrich either live cells (-/+ EIPA and -Dox conditions) or live GFP^+^ cells (+Dox condition). Approximately 100,000-150,000 cells per genotype/condition were collected and re-suspended in ∼200 μL 1X PBS containing 0.04 % (w/v) BSA (Ambion, AM2616). For library construction and sequencing, approximately 20,000 cells per genotype/condition were loaded onto a 10X Genomics Chromium Next GEM chip to generate a single cell 3’ gene expression library using the 10X Chromium Next GEM Single Cell 3’ regent kits v3.1. Pooled libraries were sequenced (paired-end, single indexing) on an Illumina NovaSeq 6000 sequencer with a standard sequencing protocol (Read 1, 28 cycles; i7 Index, 8 cycles; i5 Index, 0 cycles; Read 2, 91 cycles). The raw sequence outputs were filtered and aligned (mouse reference, mm10) to produce feature-barcode matrices using the Cell Ranger pipeline 6.1.2 on 10X Genomics Cloud Analysis.

### Bioinformatic analysis

Single-cell feature-barcode matrices were analyzed using Seurat 4.1 on RStudio server 2022.01.999. We attained an averaged number of 23789 UMIs and 17882 genes (averaged median of 2282 genes detected per cell) in the WT control (-EIPA), and 19777 UMIs and 17291 genes (averaged median of 2575 genes detected per cell) in EIPA-treated organoids from 2 separate preparations. We also obtained 22053 UMIs and 17291genes (median of 2942 genes detected per cell) in CRISPR-Cas9 control (-DOX), and 14314 UMIs and 17630 genes (median of 3507 genes detected per cell) in CRISPR-Cas9 silenced (+DOX) organoids from 1 preparation. For the pre-processing (Luecken and Theis, 2019), we selected cells based on the QC metrics (unique genes, total number of RNAs, and mitochondrial RNA) for each genotype/condition (Figure S3A). After QC filtration, we combined all datasets and then performed normalization and variance stabilization via sctransform (Hafemeister and Satija, 2019). After data integration, principal component analysis (PCA) was used to reduce the dimensionality of the combined dataset followed by constructing a shared nearest neighbor (SNN) graph (dimensions of reduction = 1:45, resolution =0.8) and building clusters with uniform manifold approximation and projection (UMAP) dimensional reduction (dimensions of reduction =1:45). To set the identities of clusters in the combined dataset, expression comparison was performed on the integrated data. In brief, markers that were distinctively enriched in each cluster were first calculated and displayed for general identification. Then, we further modified the clustering based on the average gene expression profiles of well-characterize cell-type markers. To enrich the progenitors and precursors in the secretory lineage, we subclustered a parental cluster associated with elevated secretory genes using the same PCA-SNN-UMAP workflow above (dimensions of reduction = 1:13, resolution =1), followed by subcluster identification using signature gene average expression. Lastly, we assigned each cluster with a cell-type identity in the combined dataset. A “unknown proliferative” cluster was included due to the enrichment of several minichromosome maintenance (*Mcm*) genes but without a specific cell-type identification. For downstream gene analysis, a log scale normalization (log1p scale; scale.fatctor = 10,000) and a data scaling (variables to regress out, total number of RNAs and mitochondrial RNA) were performed on the identity-assigned dataset. To quantify the relative cell number change with NHE1 inhibition, we first calculated a relative cell count of the WT control (-EIPA) and EIPA-treated clusters by calculating the number of cells in the cluster divided by the total cell count in the dataset. We then calculated a relative change by dividing the relative cell count in the EIPA-treated dataset by that of the WT control. We did not perform this calculation for the CRISPR datasets because we needed to enrich for GFP^+^ (Cas9^+^) cells to obtain the +Dox dataset, and this may have introduced a bias in the types of cells collected. To compare any gene expression between conditions (-/+EIPA or -/+Dox) in different cell types, all calculations and plotting were performed using the normalized/scaled RNA data unless stated otherwise.

### Proliferation assay and crypt cell quantification

Click-iT Edu cell proliferation kit (Invitrogen, C10340) was used to assess crypt proliferation. Day 3 organoid culture was first incubated in 10 μM Edu solution for 2 hours then proceeded for immunostaining. To quantify the proliferative cells within CD44^+^ clusters, numbers of Edu^+^ and CD44^+^ positive cells in each CD44 labeled cluster were first counted separately from confocal z-plane sectioning through the middle of a budded crypt or an unbudded organoid. For normalization, a ratio of numbers of Edu^+^; CD44^+^ double-positive cells to the numbers of total CD44^+^ cells were then calculated as a measurement of overall proliferation. To quantify other crypt cells, the numbers of *Lgr5*^+^ ISCs and *Lgr5*^-^; CD44^+^ cells (crypt progenitors) within each CD44^+^ cluster were counted and normalized to the total number of CD44^+^ cells, in the same manner as the proliferation assay. To count Paneth cells in developing organoids, organoids were fixed at day1, day2, and day3 for immunostaining. All LYZ^+^ Paneth cells present in each CD44^+^ cluster were counted in 3D confocal images and normalized to the arc length of the 3D projection of CD44^+^ clusters.

### Immunolabeling

Matrigel embedded organoids were fixed in 2% paraformaldehyde for 45 min at RT. Fixed organoids were washed 3×20 min at RT with 1X Glycine-PBS (7.5% m/v) followed by washing 2x10 min with PBS. Organoids were then permeabilized in 0.5% Triton X-100 for 15 min at RT and blocked in PBS buffer containing 0.1% BSA, 0.2% (v/v) Triton X-100, and 0.04% (v/v) Tween-20, 5∼10% (v/v) serum overnight at 4°C. Organoids were incubated overnight at 4°C with primary antibodies diluted in blocking buffer. Antibodies and dilutions included anti-CD44 (BioLegend, 103030, 1:500), anti-lysozyme (LYZ) (Dako, EC 3.2.1.17, 1:100), anti-EPHB2 (R&D, AF496, 1:25), anti-Dll1(R&D, AF5026. 1:14), and anti-Aldolase B (ALDOB; Abcam, ab75751, 1:200). After washing with blocking buffer (w/o serum) 3×1 hr at RT, organoids were incubated overnight at 4°C with fluorescent secondary antibodies (Molecular Probes) diluted 1:200 in blocking buffer. After incubation, organoids were washed 3×1 hr with blocking buffer (w/o serum) and then 20 min with PBS. Washed organoids were stained with Hoechst 33342 (Molecular probes, 1:1000) and stored in PBS at 4°C. Fluorescence images were acquired using a customized spinning disk confocal (Yokogawa CSU-X1) on a Nikon Ti-E microscope with a 40X water objective equipped with a Photometrics cMYO cooled CCD camera.

For image analysis, background of the region of interest in each section of Z-stacks was corrected by subtracting a value of mean intensity from a sample-free region and single optical sections containing representative intestinal crypts were chosen for quantification. Individual cells were segmented by the membrane dye. To identify major cell types in the crypt, Paneth cells were selected based on size and presence of visible intracellular granules under both bright-field and mCherry channels, ISCs were identified based on their location which is intermingled between two Paneth cells, other cells along the crypt column starting at a position two-cell above the upmost Paneth cell and ending at the crypt neck, the upmost part of a budded crypt, were further specified as the crypt column and the crypt neck. In the crypt without a bud, up to four of the cells closest to the upmost Paneth cell were chosen as the column for pHi measurement.

Because the crypt region and villus region were less distinguishable by live-cell imaging in the day 1 organoid during development, the column region was not specified for pHi determination on day 1. To calculate mCherry/SEpHluorin intensity ratios, fluorescence intensity was measured in the selected individual Paneth cells, ISCs, and cells located in the column and neck using ImageJ. In Microsoft Excel, the pHi values were determined by applying those intensity ratios to a standard curve generated by graphing the nigericin pH values (Y-axis) against their dependent intensity ratios (X-axis). Because of the mCherry/SEpHluorin intensity ratio in the cells in the organoid villus (above the crypt neck) was too high to be calibrated, the pHi in those cells were not determined.

### Single cell reassociation assay

NHE1 silencing was induced by adding 2 μg/mL Dox to the inducible-NHE1 CRISPR Cas9-GFP organoids culture for 2 days. On the day of sorting, the NHE1 CRISPR silenced organoids and *Lgr5*^DTR-GFP^ organoids were first dissociated and then stained for CD44 (BioLegend, 103030, 1:250), CD24 (BD, 553262, 1:250), and Live/Dead (Invitrogen, L23105, 1:1000) in the sorting buffer. FACS was performed on the BD FACSAira. Live *Lgr5*^+^ ISCs (*Lgr5*-DTR-GFP^high^, CD44^high^, CD24^low^, and side-scatter^low^) were sorted and mixed with either the sorted live WT Paneth cells (*Lgr5*-DTR-GFP^negative^, CD44^high^, CD24^high^, and side-scatter^high^) or the sorted live NHE1-silenced Paneth cells (Cas9-GFP^high^, CD44^high^, CD24^high^, and side-scatter^high^) at 1:1 ratio (400-500 cells each) as previously described^30^. In addition, groups containing the *Lgr5*^+^ ISCs alone, the *Lgr5*^+^ ISCs + WNT3A (20% v/v WNT3A in single-cell growth media), the WT Paneth cells alone, and the NHE1-silenced Paneth cells alone were also included as controls for normalization. All groups were grown in single cell growth media for 3 days and then switched to normal ENR media. To evaluate organoid formation ability from the ISC-Paneth cell pairs, the number of organoids formed in all the groups (if any) was counted on day 8 and normalized to the *Lgr5*^+^ ISCs + WNT3A group.

### Lineage tracing

In general, lineage tracing in the *Atoh1*^CreERT2^; *Rosa26*^tdTomato^ and in the *Lgr5*^CreER^; *Rosa*26^RFP^organoids were prepared by treating the day 1 organoids with 0.1 μM 4-hydroxytamoxifen (TAM; Sigma-Aldrich, 579002) for 26 hours followed by washing and passaging. Reseeded organoids were cultured with vehicle control or 5 μM EIPA for 2-3 days either in a standard culture incubator before immunostaining for DLL1 and ALDOB, or in a live-cell imaging chamber for confocal microscopy for Paneth cell production, recorded every 24 hr, with z-stacks of 2 μm optical sections covering the entire crypt region. To reveal overall ATOH1^+^ secretory cell production, day 1 control and EIPA-treated *Atoh1*^CreERT2^; *Rosa26*^tdTomato^ organoid culture were added with 0.1 μM TAM for 24 hours first before starting a 48 hr time-lapse imaging (day 2 to day 3), which were acquired at 30-min intervals, in a 2-μm z step size per optical section. To quantify lineage tracing, numbers of RFP^+^ or tdTomato^+^ cells were counted in the crypt regions, indicated by immunolabelled CD44^+^ clusters in the *Lgr5*^CreER^; *Rosa26*^RFP^ organoids, or by clusters of live ATOH1^+^ cells and Paneth cells in the *Atoh1^CreERT2^*; *Rosa26*^tdTomato^ organoids. To assess the Paneth cell production, changes of newly generated ATOH1^+^ Paneth cells (RFP^+^/dense granule^+^) numbers within 24 hrs were normalized to the number of total ATOH1^+^ cells (RFP^+^) as well as the Atoh1^+^ non-Paneth cells (RFP^+^/dense granule^-^). Similarly, for measuring the overall secretory cell production, changes of newly produced ATOH1^+^ cells (RFP^+^) numbers from day2-3 were normalized to the number of ATOH1^+^ cells seen on day 2. To determine differentiation capability of *Lgr5*^+^ ISCs into DLL1^+^ enriched secretory progenitors, the Paneth cells, and the enterocytes, percent of crypts containing RFP^+^/CD44^+^/DLL1^+^ (secretory progenitors), RFP^+^/CD44^+^/LYZ^+^(Paneth cells), and organoids containing RFP^+^/CD44^-^/ALDOB^+^ cells were calculated. The number of RFP^+^/CD44^+^/DLL1^+^, RFP^+^/CD44^+^/LYZ^+^, and RFP^+^/CD44^-^/ALDOB^+^ cells were also quantified in each crypt or organoid as a measurement for *Lgr5*^+^ ISC differentiation.

### RNA extraction and quantitative reverse transcription PCR (RT-qPCR)

Total RNA was extracted from organoids using the RNeasy plus mini kit (Qiagen, 74134). RNA was used to generate cDNA using the iScript cDNA synthesis kit (Bio-Rad, 1708890). The mRNA level of target genes was accessed by RT-qPCR on a Real-time PCR system (Applied Biosystems) using the SYBR green supermix reaction (Bio-Rad, 170-8882). The relative expression of targets was analyzed using the double delta C_t_ method by normalizing to *Gapdh*. Primer sequences were below: *Lgr5* forward (5’-GGGAGCGTTCACGGGCCTTC-3’), *Lgr5* reverse (5’-GGTTGGCATCTAGGCGCAGGG-3’); *Lyz* forward (5’-GGAATGGATGGCTACCGTGG-3’), *Lyz* reverse (5’-CATGCCACCCATGCTCGAAT-3’); *Alpi1* forward (5’-AGGATCCATCTGTCCTTTGG-3’), *Alpi1* reverse (5’-ACGTTGTATGTCTTGGACAG-3’); *ChgA* forward (5’-CTCGTCCACTCTTTCCGCAC-3’), *ChgA* reverse (5’-CTGGGTTTGGACAGCGAGTC-3’); *Muc2* forward (5’-ATGCCCACCTCCTCAAAGAC-3’), *Muc2* reverse (5’-GTAGTTTCCGTTGGAACAGTGAA-3’); *Hes1* forward (5’-GCTCACTTCGGACTCCATGTG-3’), *Hes1* reverse (5’-GCTAGGGACTTTACGGGTAGCA-3’); *Atoh1* forward (5’-GCCTTGCCGGACTCGCTTCTC-3’), *Atoh1* reverse (5’-TCTGTGCCATCATCGCTGTTAGGG-3’); *Axin2* forward (5’-AGTGTCTCTACCTCATTTTCCG-3’), *Axin2* reverse (5’-CTTTCCAGCTCCAGTTTCAGT-3’); *Ascl2* forward (5’-GCTGCTTGACTTTTCCAGTTG-3’), *Ascl2* reverse (5’-CACTAGACA GCATGGGTAAGG-3’); *Olfm4* forward (5’-TGAAGGAGATGCAAAAACTGG-3’), *Olfm4* reverse (5’-CTCCAGCTTCTCTACCAAGAGG-3’); *Gapdh* forward (5’-TGTGTCCGTCGTGGATCTGA-3’), *Gapdh* reverse (5’-CCTGCTTCACCACCTTCTTGA-3’).

### Statistics

Statistical analysis was performed using GraphPad Prism 8. Outliers were determined using the ROUT method by fitting data with nonlinear regression and were excluded from other statistical tests. Data fitting a normal distribution were analyzed using a standard t-test/Welch’s t-test. The non-normally distributed data were analyzed by the Mann-Whitney ranking test.

---

**Figure S1.**
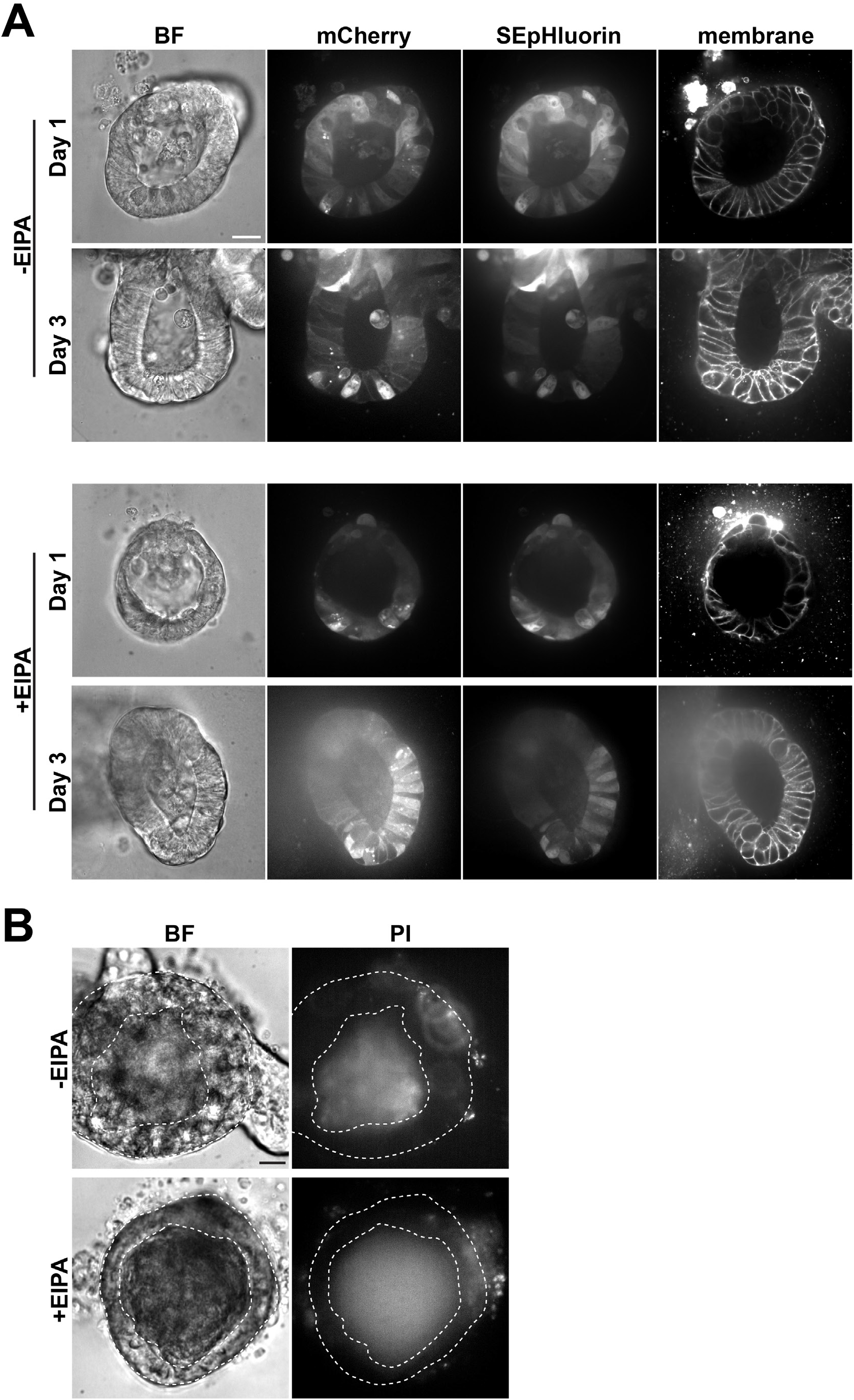
mCherry-SEpHluroin and cell viability in organoids. (**A**) Representative live-cell images of mCherry-SEpHluorin in the crypt region in day 1 and day 3 organoids in the absence (Control) and presence of 5 μM EIPA. Panels show brightfield (BF) individual and merged mCherry and SEpHluorin signal, and far-red membrane dye to determine individual cells. **b,** Propidium iodide (PI) staining as an index of cell viability in day 3 organoids in the absence and presence of EIPA. Space between dashed circles indicates epithelial layer. Inner dashed circle indicates the lumen containing apoptotic cells with positive staining. All scale bars, 20 mm.

**Figure S2.**
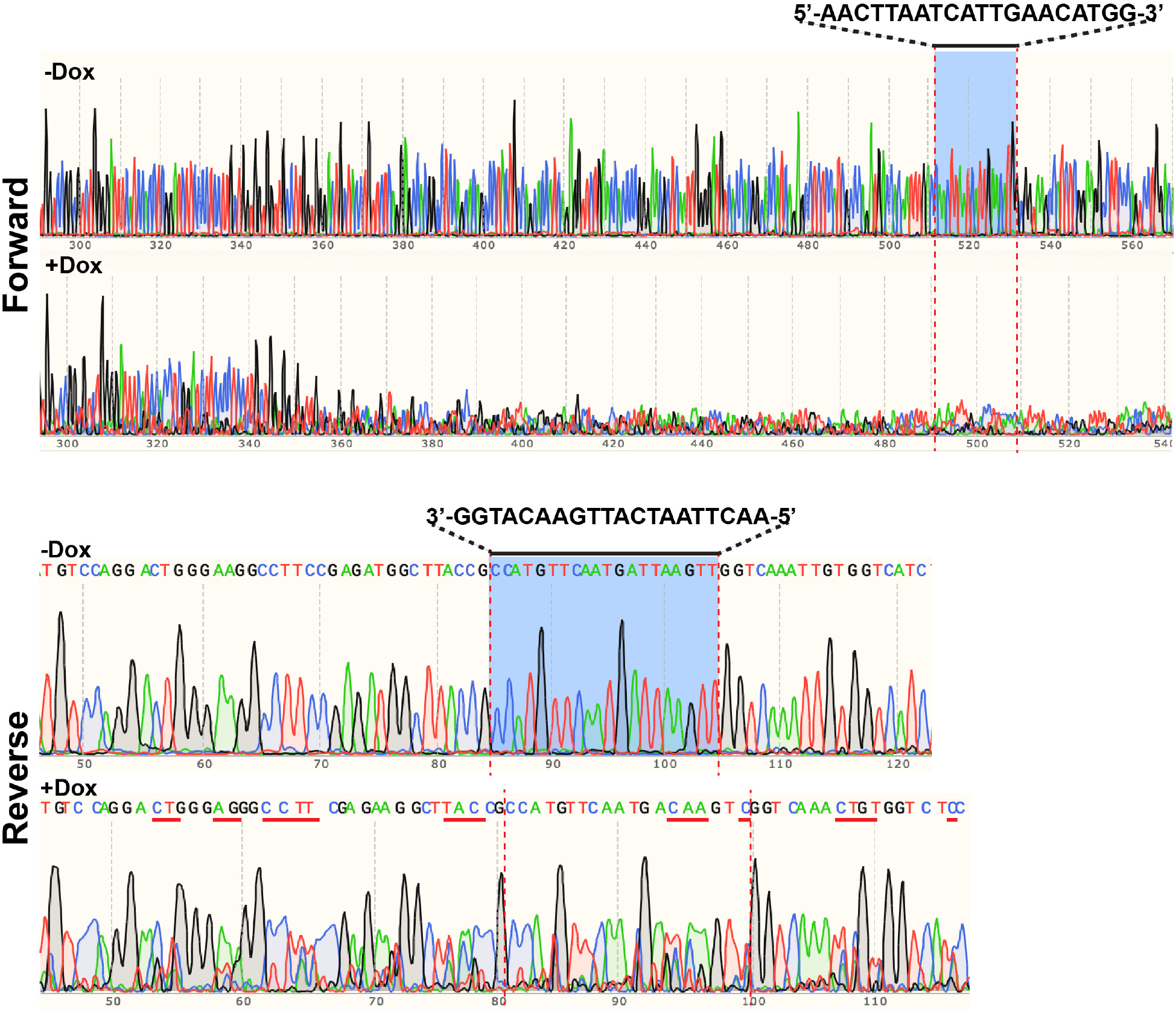
Comparison of Sanger sequencing of PCR fragments of NHE1 gene from non-induced (-Dox) and induced (+Dox) organoids. Results are generated using forward and reverse sequencing primers, respectively. The Cas9 guide RNA targeting region is indicated by a black line with red dashed lines. Reduction of signal in forwarding sequencing indicates large disruption of DNA sequence. A red solid line in reverse sequencing indicates CRISPR indels.

**Figure S3.**
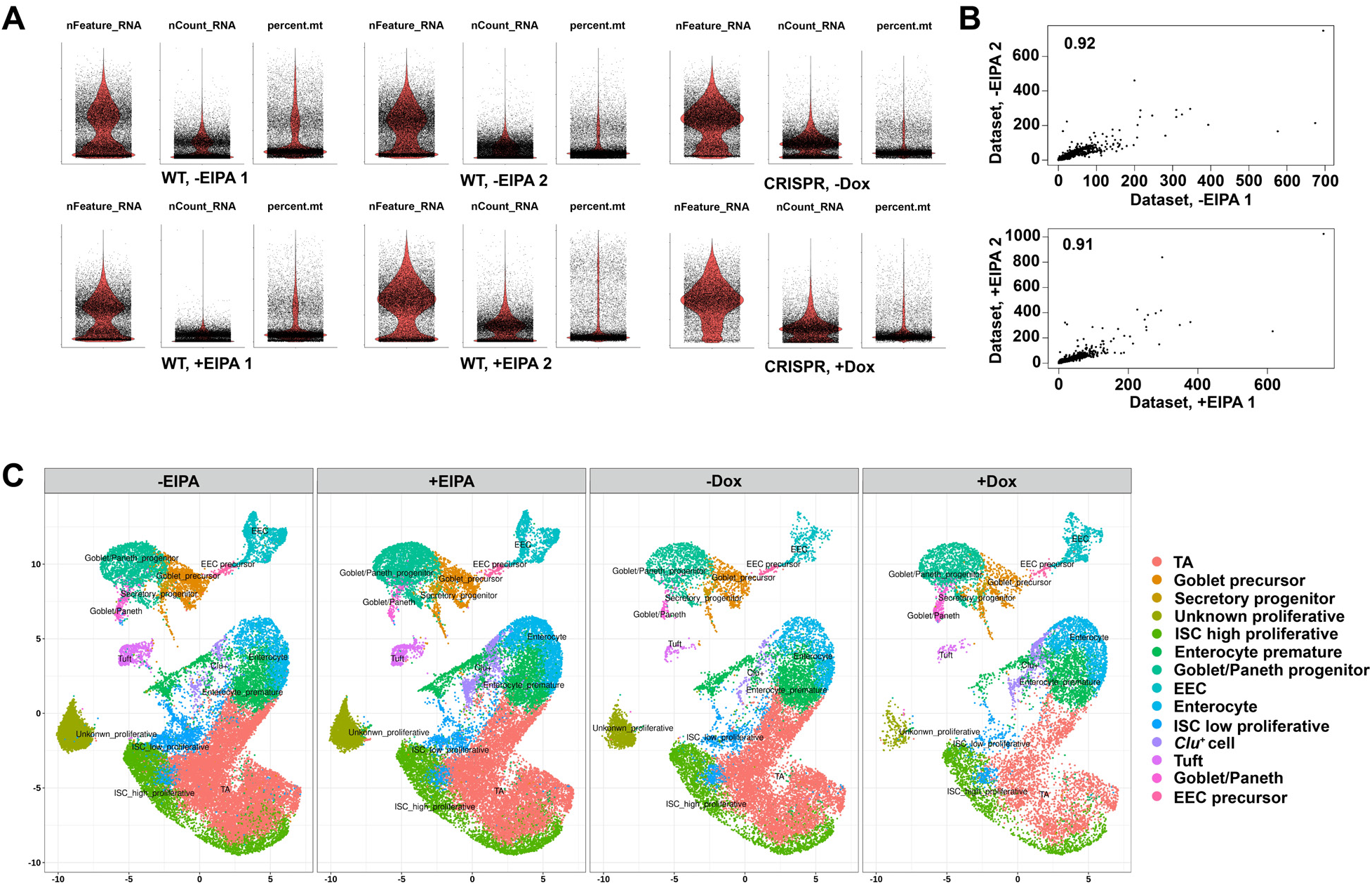
Quality control of scRNA-seq. (**A**) Visualization of the quality control metrics of datasets (Methods). Violin plots showing nFeature_RNA (unique genes), nCount_RNA (total number of RNAs), and percent.Mt (mitochondrial RNA, low quality and dying cells). (**B**) Scatter plots showing the relationship between biological replicates of organoids preparation (Methods). The Pearson correlation coefficient, indicated in the upper left of each graph, is calculated using the average expression profiles of individual datasets. (**C**) UMAP visualization of cell-identity clusters sorted by the conditions, WT+/-EIPA, CRISPR+/-Dox.

**Figure S4.**
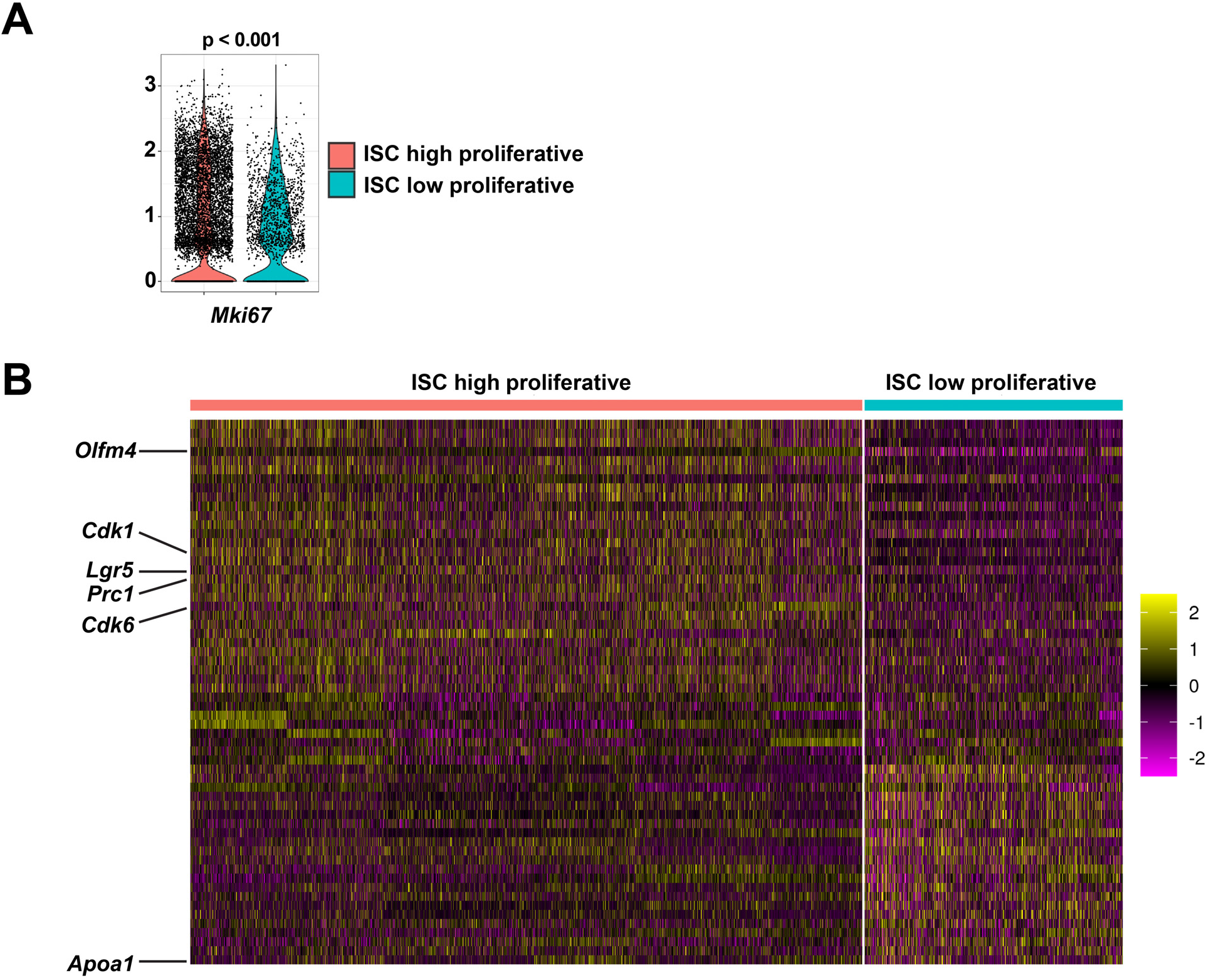
Identification of high proliferative and low proliferative ISC subtypes. (**A**)Violin plot showing the expression level of *Mki67* between the distinct ISC subclusters. Data are assessed using the Student-t test. (**B**) Heatmap showing signatures of high proliferative and low proliferative ISC subtypes. Colored expression level is relative to the mean expression of cell population, 0 (population mean) ± 2 (SD). Cell cycle signature genes (*Cdk1*, *Prc1*, and *Cdk6*), Stem cell-specific genes (*Lgr5* and *Olfm4*) and the absorptive-associated gene (*Apoa1*) are highlighted (left row).

**Figure S5.**
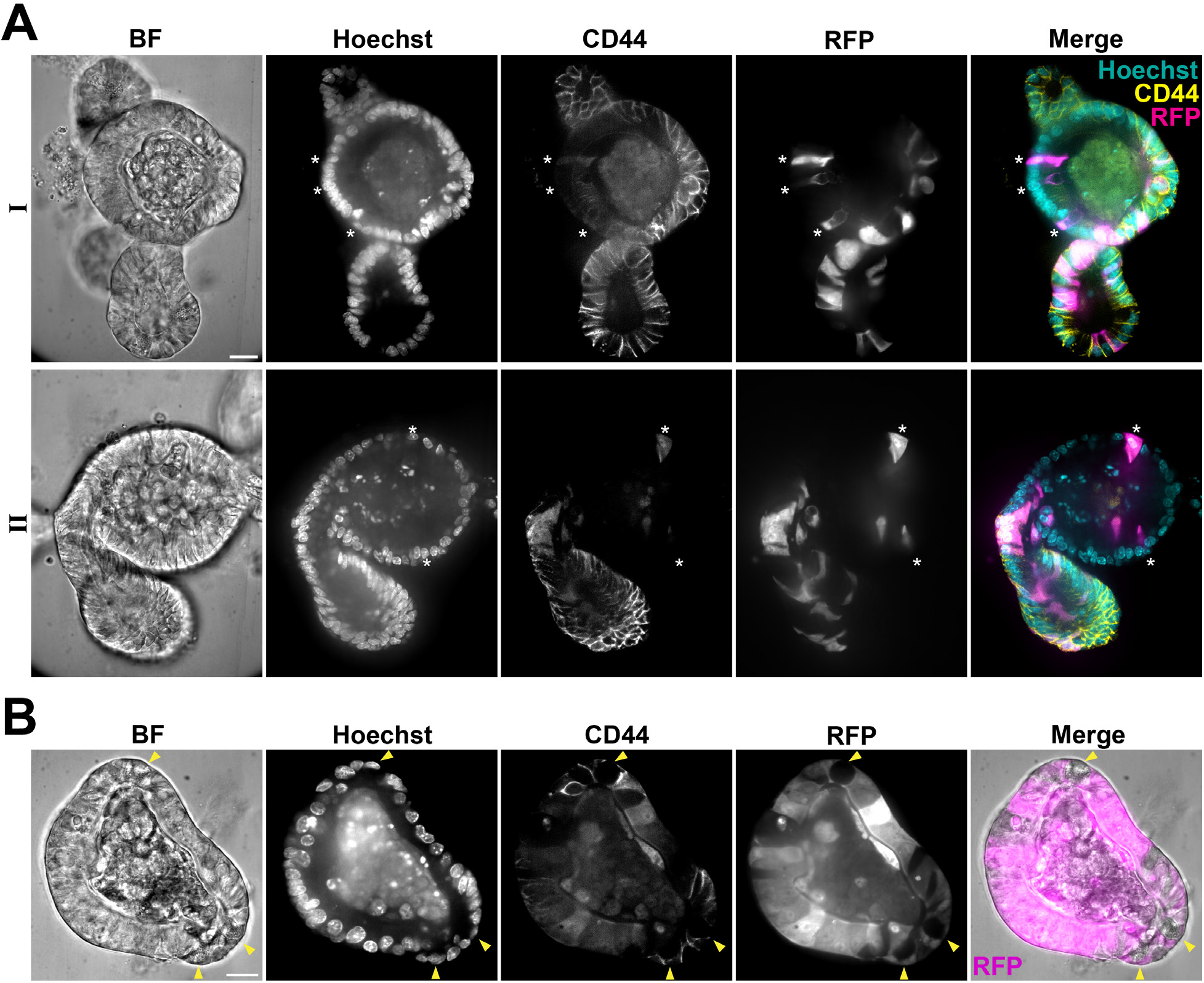
Lineage tracing of crypt and villus cells in *Lgr5*^CreER^;*Rosa26*^RFP^ organoids. (**A**) Representative images show the presence of labeled *Lgr5*^+^ ISC progeny (RFP^+^) in crypt (CD44^+^) and villus (CD44^-^) regions of day 3 organoids. Organoids are treated with 4-hydroxytamoxifen on day 1 followed by washing and reseeding on day 2. Immunolabeled images show 2 examples (I, II) of day 3 reseeded organoids. (**B**) Example of an EIPA-treated organoid with robust labeling in the *Lgr5*^+^ ISC progeny. Arrowhead indicates Paneth cells, which are identified by visible dense granules and large cell size. All scale bars, 20 μm.

**Figure S6.**
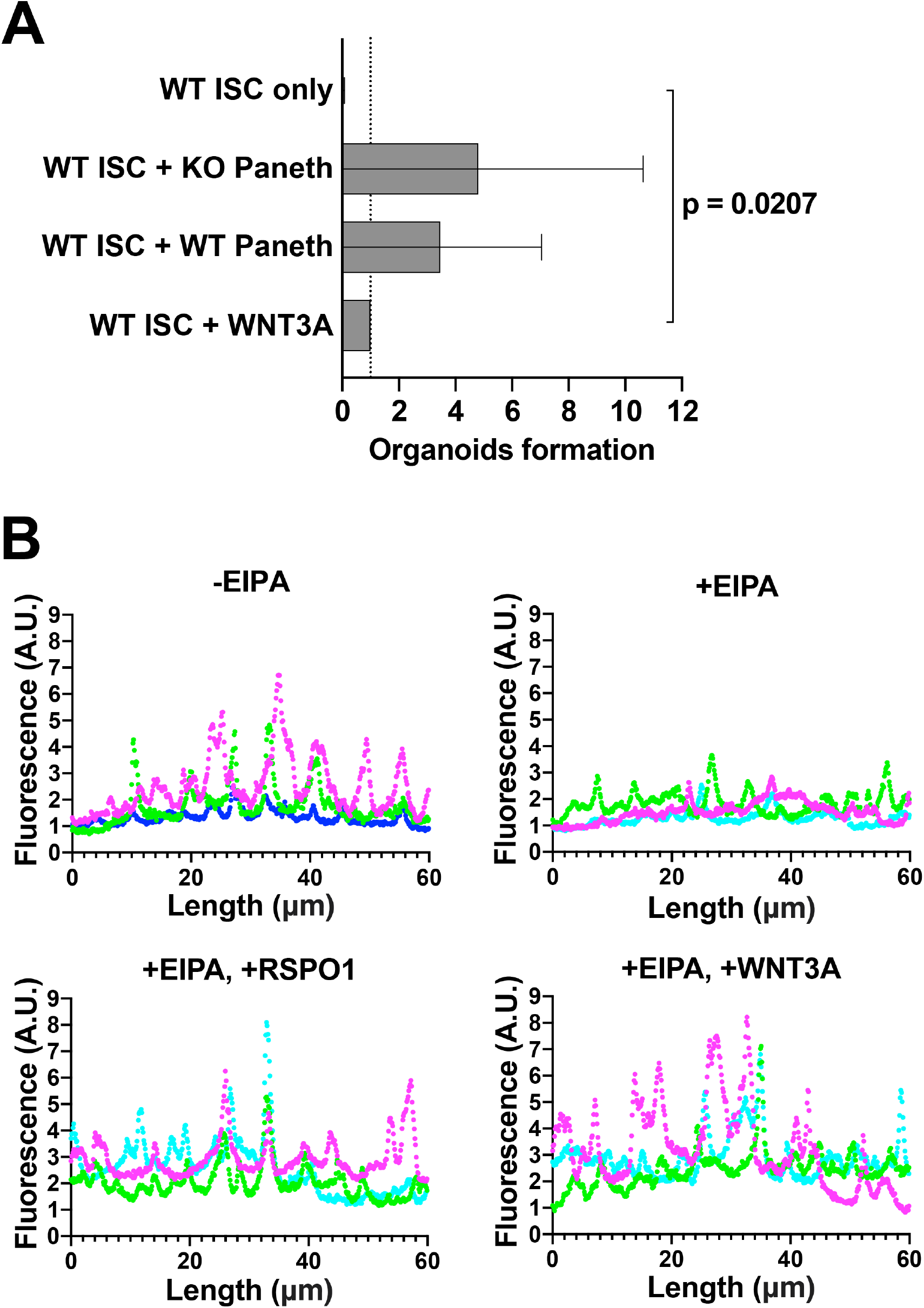
Quantification of organoids formation and EPHB2 staining. (**A**) Relative efficiency of *Lgr5*^+^ ISC-Paneth cell single cell reassociation, determined by the number of organoids formed. Data are normalized to the positive control (single *Lgr5*^+^ ISCs alone with WNT3A) and show the means of 2 independent preparations, with statistical analysis by the Wilcoxon test. Data with a statistically significant difference are specified with a p value, otherwise are not significantly different. (**B**) Representative line plots of intensity of EPHB2 immunolabeling in crypt region of day 3 organoids with and without WNT rescue. Fluorescence intensity is determined by drawing a line across lateral layers of crypt cells using the line toolfunction in ImageJ. Each plot shows 3 individual crypt regions indicated by different colors (n=3).

**Figure S7.**
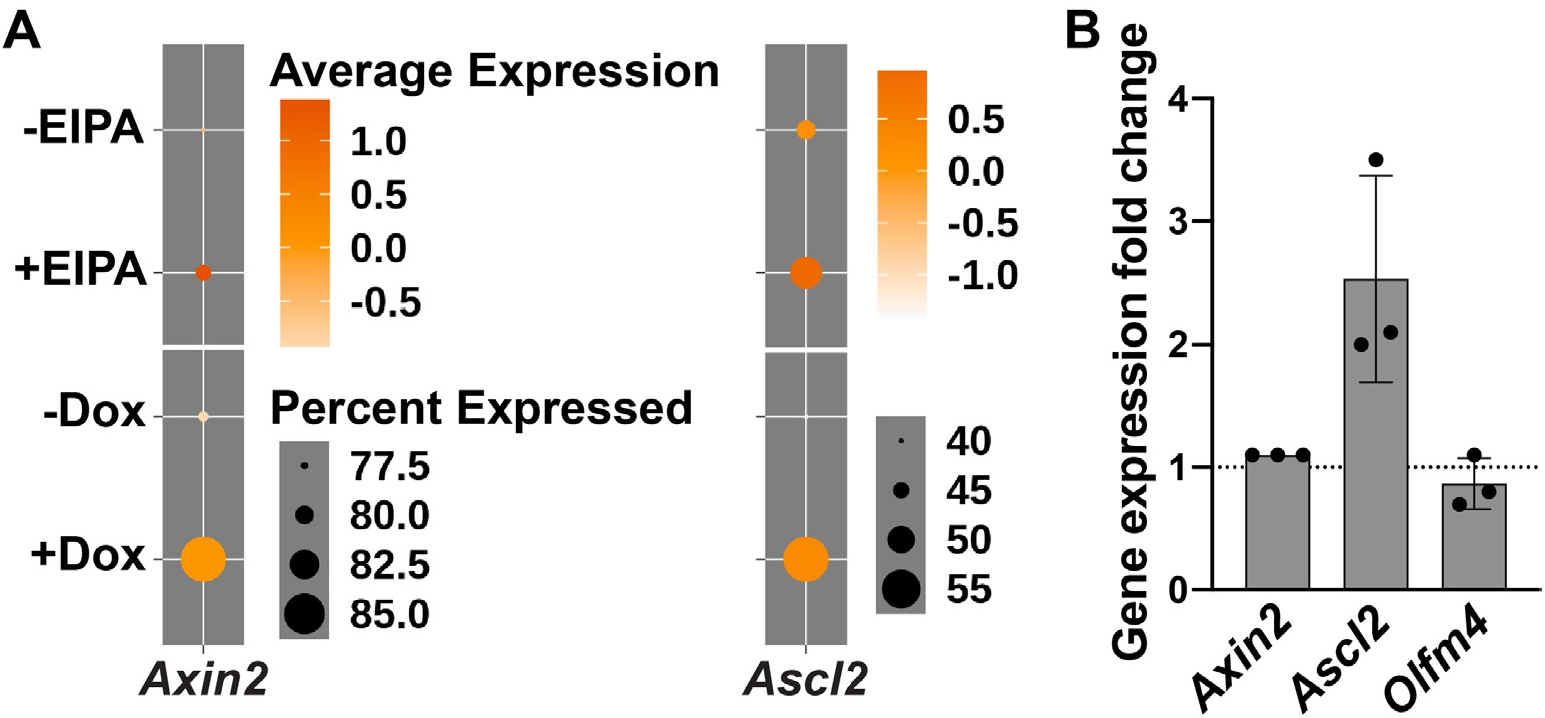
Expression of WNT responsive and non-WNT responsive genes with NHE1 inhibition. (**A**) Dot plots showing the average expression level of WNT target genes, *Axin2* and *Axcl2* in the *Lgr5*^+^ ISC cluster. All dot plots are colored by average expression, 0 (population mean) ± SD. (**B**) Expression (mean ± SD) of *Axin2*, *Ascl2* and *Olfm4* in EIPA-treated organoids relative to non-treated organoids (controls, n=3).

## Video S1

Video of mCherry-SEpHluroin fluorescence ratio in a budding crypt (day 1 to day 2), corresponding to **Figure 1D**. Fluorescence ratios are calibrated to pH values using nigericin-containing buffers to show that a dynamic pHi gradient, lower at the crypt base and higher at the crypt neck, is generated during crypt budding. Left view, bright field. Right view, ratio (mCherry/SEpHluorin).

## Video S2A

Video of crypt budding in an untreated control *Lgr5*^DTR-GFP^ organoid from day 0 to day 3, corresponding to **Figure 2A**. The crypt region elongates and forms a budded crypt. Left view, bright field. Right view, *Lgr5*^+^ ISC.

## Video S2B

Video of impaired crypt budding in an EIPA-treated *Lgr5*^DTR-GFP^ organoid from day 0 to day 3, corresponding to **Figure 2A**. The crypt region initiates a protrusion but fails to form a budded crypt. Left view, bright field. Right view, *Lgr5*^+^ ISC.

## Video S3A

Video shows the production of secretory cells in the crypt region of an untreated control *Atoh1*^CreERT2^;*Rosa26*^tdTomato^ organoid from day 2 to day 3, corresponding to **Figure 6F**. Newly produced ATOH1^+^ (tdTomato^+^) secretory cells appear in the crypt as the crypt elongates and forms a bud.

## Video S3B

Video shows attenuated production of secretory cells in the crypt region of an EIPA-treated *Atoh1*^CreERT2^;*Rosa26*^tdTomato^ organoid from day 2 to day 3, corresponding to **Figure 6F**. No evident increase of newly produced ATOH1^+^(tdTomato^+^) secretory cells appears in the unbudded crypt region.

